# Regulation of the expression of *nucS*, a key component of mismatch repair system in mycobacteria

**DOI:** 10.1101/2025.07.23.666449

**Authors:** Esmeralda Cebrián-Sastre, Ángel Ruiz-Enamorado, Alfredo Castañeda-García, Susanne Gola, Pablo García-Bravo, Leonor Kremer, Jesús Blázquez

## Abstract

Alterations in the expression of the mismatch repair (MMR) system can lead to transient hypermutation, accelerating the emergence of adaptive mutations under stress conditions, such as antibiotic pressure. While most bacteria and eukaryotes rely on the MutS and MutL protein families for MMR, many archaeal and actinobacterial species, including the major human pathogen *Mycobacterium tuberculosis*, lack these components. Instead, they utilize NucS, a structurally and evolutionarily distinct enzyme driving a non-canonical MMR pathway. Given the role of MMR in mutation control, understanding how *nucS* expression is regulated could be essential for uncovering the molecular basis of antibiotic resistance development in mycobacteria.

In this study, we defined the promoter region and transcription start site of the *nucS* gene in *Mycobacterium smegmatis*. We found that *nucS* expression is growth phase-dependent in both *M. smegmatis* and *M. tuberculosis*, significantly decreasing during the stationary phase. This downregulation mirrors that of canonical MMR components, aligning with replication activity. Additionally, we present evidence that the alternative sigma factor σ^B^ may negatively regulate *nucS* expression during the stationary phase.

We also identified candidate compounds capable of modulating *nucS* expression, underscoring its responsiveness to environmental cues. These findings deepen our understanding of mycobacterial stress adaptation and provide a groundwork for further investigation into the molecular mechanisms underlying antibiotic resistance.

Strikingly, our work reveals a case of double convergent evolution: both canonical (MutS/MutL) and non-canonical (NucS) pathways have independently evolved not only the same DNA repair function, but also similar regulatory frameworks for genome integrity preservation under stress conditions.

**IMPORTANCE:** Bacteria like *Mycobacterium tuberculosis*, which causes tuberculosis, develop antibiotic resistance through mutations. Most organisms use a MutS/MutL-based system called mismatch repair to correct DNA replication errors and prevent harmful mutations. However, some bacteria, including mycobacteria, rely on a unique repair protein called NucS. This study explores how the gene encoding NucS is regulated in mycobacteria.

We found that less NucS is produced when bacteria stop growing, which mirrors what happens with other DNA repair systems. We also discovered that a regulatory protein called SigB may help reduce its production during this phase. In addition, we identified chemical compounds that influence how much NucS bacteria produce, showing that its levels respond to environmental conditions.

Importantly, the study reveals that different DNA repair systems—despite being unrelated—have evolved similar ways to protect bacteria under stress. These findings help scientists understand how mycobacteria adapt and could aid in fighting antibiotic resistance.

## INTRODUCTION

Bacteria continuously face fluctuating environmental conditions that demand rapid and coordinated stress responses. These adaptive strategies frequently involve a transient elevation of mutation rates, thereby increasing genetic variability and enhancing survival under specific adverse conditions (1).

In *Escherichia coli* and related microorganisms, nutrient limitation during the stationary phase triggers a general stress response that can increase the mutation rate either by inducing the expression of the error-prone DNA polymerase (Pol IV) or by repressing the expression of key components of the mismatch repair (MMR) system (1). Moreover, since MMR system constrains recombination between non-identical (homeologous) sequences, its downregulation in the stationary phase may further facilitate the exchange of divergent alleles, broadening the adaptive landscape available to the cell (1).

Detailed investigation in *E. coli* have shown that levels of MMR proteins MutS and MutH are reduced by approximately four-fold and two-fold, respectively, during the stationary phase compared to the exponential growth (2), whereas MutL levels remain constant across both phases (2, 3). Even though MutL levels appear stable, evidence suggests that its active concentration becomes limiting during the stationary phase, contrasting with the proportional decline in MutS and MutH, which likely mirrors the reduced replication demand (2, 4, 5).

Global regulators such as the alternative sigma factor RpoS (σ^S^) and the RNA chaperone Hfq play essential roles in adjusting MutS and MutH protein levels during stress (2). In the exponential phase, Hfq destabilizes *mutS* transcripts via an RpoS-independent mechanism. As cells transition to the stationary phase, Hfq further downregulate MutS via both RpoS-dependent and independent pathways, potentially through intermediaries like RNases, proteases, or other RpoS-controlled factors (2). Hfq and RpoS also seem to regulate MutH levels through the same pathway during the stationary phase (2). In addition, subinhibitory concentrations of β-lactam antibiotics induce the *rpoS* regulon. Under these conditions, the RpoS-controlled regulatory sRNA SdsR targets *mutS* mRNA, preventing its translation and consequently reducing MMR activity (6). Recent evidence also implicates an RNA G-quadruplex structure, formed by guanine-rich sequences within the *mutS* coding region, as a potential regulatory element (7).

While the canonical MMR proteins MutS and MutL are conserved across all domains of life, from bacteria to humans, many Archaea and nearly all member of the phylum Actinobacteria, including major pathogens such as *Mycobacterium tuberculosis*, lack these components (8–10). Remarkably, these microorganisms maintain low mutation rates, suggesting the existence of an alternative MMR pathway. This observation led our research group to identify a non-canonical MMR system in Actinobacteria, based on a homolog of the archaeal endonuclease NucS (10). The in vivo role of NucS as an MMR protein was first demonstrated in the nonpathogenic species *Mycobacterium smegmatis*, where its disruption leads to a hypermutator phenotype, a bias towards transition mutations, and increased homeologous recombination, hallmark features of MMR deficiency (10, 11). These findings underscored the essential role of NucS in maintaining genome integrity in Actinobacteria (10, 11).

Despite these advances, the molecular mechanisms regulating *nucS* expression remain unknown. We hypothesized that *nucS* expression is subject to transcriptional and/or post-transcriptional regulation, influenced by the physiological state of the cells, environmental stressors (including certain antibiotics), and host factors such as oxidative and nitrosative stress. Indeed, recent evidence has suggested that nitrosative agents may influence *nucS* expression (12). Unraveling these regulatory mechanisms could prove invaluable for preventing resistance development and optimizing antibiotic therapies.

In this work, we investigated the expression profile of mycobacterial *nucS* across growth phases and examined the potential regulatory role of the sigma factor σ^B^, a functional analog of *rpoS* (13–16), after identifying putative σ^B^ binding boxes in the *nucS* promoter region. Furthermore, to explore the influence of environmental factors, we evaluated the impact of a number of chemical compounds on *nucS* expression using a *nucS*::*gfp* transcriptional fusion. The dynamic regulation of *nucS* expression may induce transient hypermutation, potentially increasing genetic variability and providing an adaptive advantage under certain stressful conditions.

## RESULTS

### Determination of the Transcription Start Site of *nucS* in *M. smegmatis*

To initiate the study of *nucS* expression, we determined its transcription start site (TSS) using the 5’-RACE technique. A TSS was identified that coincides with the gene’s translation start site, consistent with findings from the global analysis by Martini *et al.* (17) (**Figure 1A**). This overlap between the transcription and translation start sites, resulting in the absence of a 5’-UTR, is commonly observed in mycobacteria (18).

**Figure 1.**
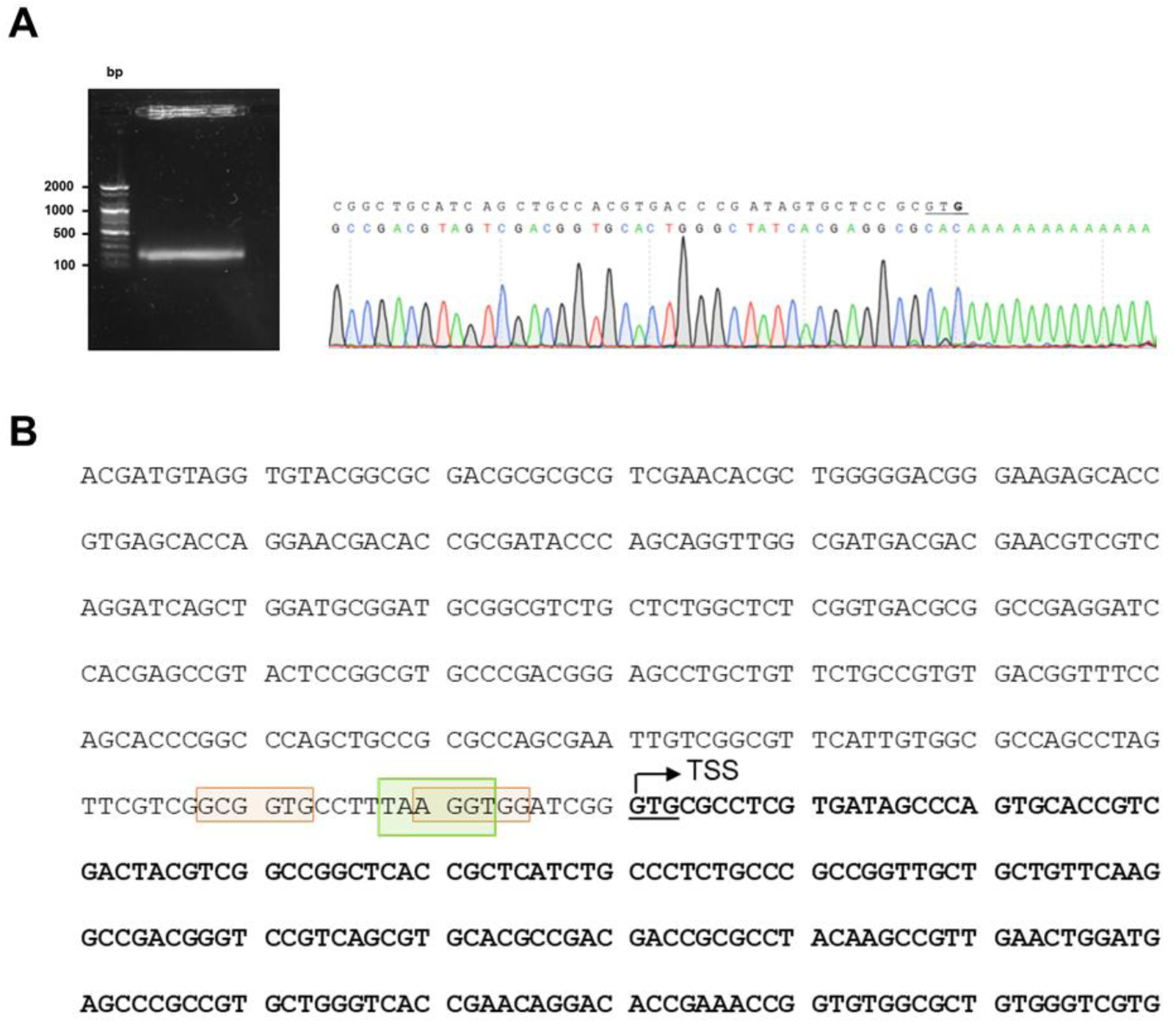
Identification of *nucS* TSS. A. Results of 5’-RACE experiment. A gel image showing the band obtained after performing the 5’-RACE technique, which was sequenced to identify the TSS of *nucS* in *M. smegmatis*. The TSS of *nucS* is highlighted in bold, corresponding to the complementary base to the first nucleotide following the poly-A tail. The TSS coincides with the translation start of *nucS* (the first G of the start codon GTG, which is underlined). **B. Upstream sequence of *nucS* in *M. smegmatis*.** The start of the coding sequence of the *nucS* gene is shown in bold, with the GTG start codon underlined. The identified transcription start site (TSS), determined by 5’-RACE, is indicated with an arrow. The colored boxes represent potential –10 regions for binding of sigma factors in *M. smegmatis*: green for σ^A^ (TANNNT) (18) and orange for σ^B^ (NNGNNG) (19).

The determination of the TSS of *nucS* allows us to better locate the promoter region of the gene. Close to the TSS, we identified the consensus sequence of the –10 box for σ^A^ factor binding (TANNNT), homologous to σ^70^ of *E. coli* and commonly found in constitutively expressed genes (18) (**Figure 1B**). We did not identify the consensus sequence for the –35 box for σ^A^ binding, as the –35 box is rarely conserved in mycobacteria (19). Based on the consensus sequences reported by Newton-Foot and Gey van Pittius (19), potential –10 boxes for other sigma factors were detected near the σ^A^ binding site. While these sequences are less conserved than those of σ^A^, we identified two putative consensus sequences for the –10 box of σ^B^ (NNGNNG), a sigma factor that plays a crucial role in response to various stress conditions and adaptation to the stationary phase (**Figure 1B**).

### Construction of GFP reporter strains for the study of *nucS* expression

In this study, reporter strains were constructed to analyze the expression levels and regulatory mechanisms of the *nucS* gene. Specifically, three transcriptional reporter fusions were constructed, each containing *nucS* upstream regions of varying lengths followed by the GFP gene. The first fusion, referred to as the “short fusion,” included a 73 bp region directly upstream of the *nucS* start codon (pSGV53-P*_nucS_*-73-*gfp*), which may correspond to the core promoter region according to the identified TSS and the predicted sigma factor binding sites. The second fusion, referred to as the “long fusion,” encompassed a 408 bp region upstream of the *nucS* start codon (pSGV53-P*_nucS_*-408-*gfp*), including the same region plus additional upstream sequences that may contain regulatory elements. The third fusion spanned the same 408 bp region but excluded the first 46 bp (pSGV53-P*_nucS_*-408Δ46-*gfp*), which corresponds to the intergenic sequence immediately upstream of *nucS*, and therefore likely lacked the core promoter region (**Figure 2**).

**Figure 2.**
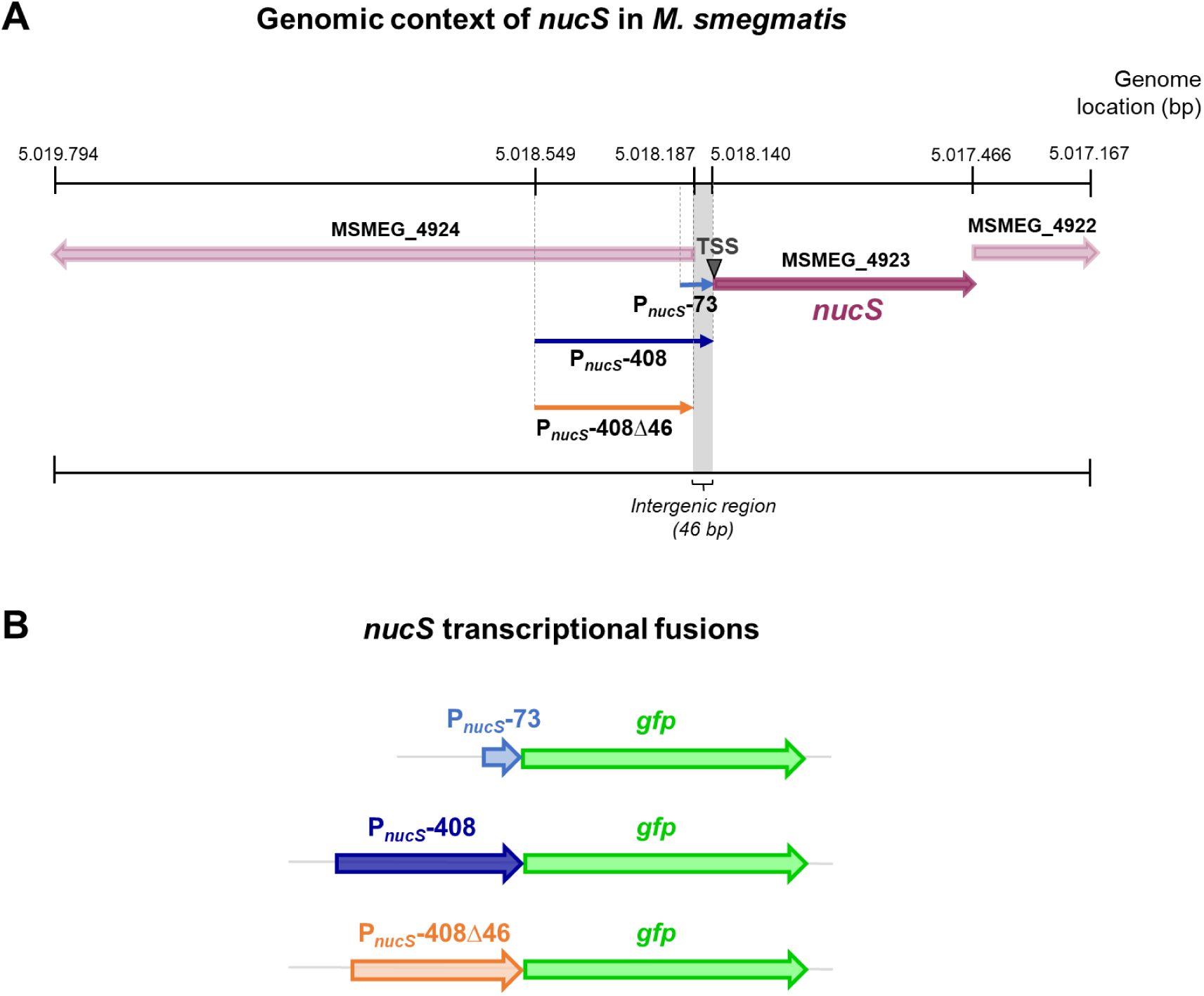
Genomic context of *nucS* in *M. smegmatis* and schematic of *nucS* transcriptional fusions. A. Genomic context of the *nucS* gene in *M. smegmatis* mc^2^ 155. The position of *nucS* in the genome of *M. smegmatis* mc^2^ 155 (NC_008596) is shown, along with the upstream regions of the gene (P*_nucS_*-73, P *_nucS_*-408 and P *_nucS_*-408Δ46) that were included in the transcriptional fusions with *gfp*. The position of the *nucS* TSS is also indicated. B. Schematic representation of the *nucS::gfp* transcriptional fusions. The three *nucS* upstream regions of different lengths cloned into the reporter vectors are shown, each followed by the *gfp* gene.

The plasmid pSGV53 served as the backbone for constructing the reporter fusions (**Table S1**). This plasmid is a replicative vector with a replication origin derived from pAL5000, maintaining approximately 5 copies per cell (20, 21), and originally carries the gene for the eGFP protein under the control of the constitutive promoter P*_mpt64_* (22). To generate the reporter plasmids, the constitutive promoter was replaced with the upstream regions of *nucS*, enabling the fluorescence signal to reflect the activity of these regulatory regions.

### *nucS* expression is growth phase-dependent

To investigate *nucS* expression, the reporter plasmids were transformed into the wild – type *M. smegmatis* strain, and fluorescence levels during growth were monitored using a spectrofluorometer. Fluorescence intensity (FI) was normalized to OD_595_ values at each time point (**Figure 3A**). No differences in growth were observed between the strains carrying the reporter vectors and the non-transformed control strain (**Figure S1**).

**Figure 3.**
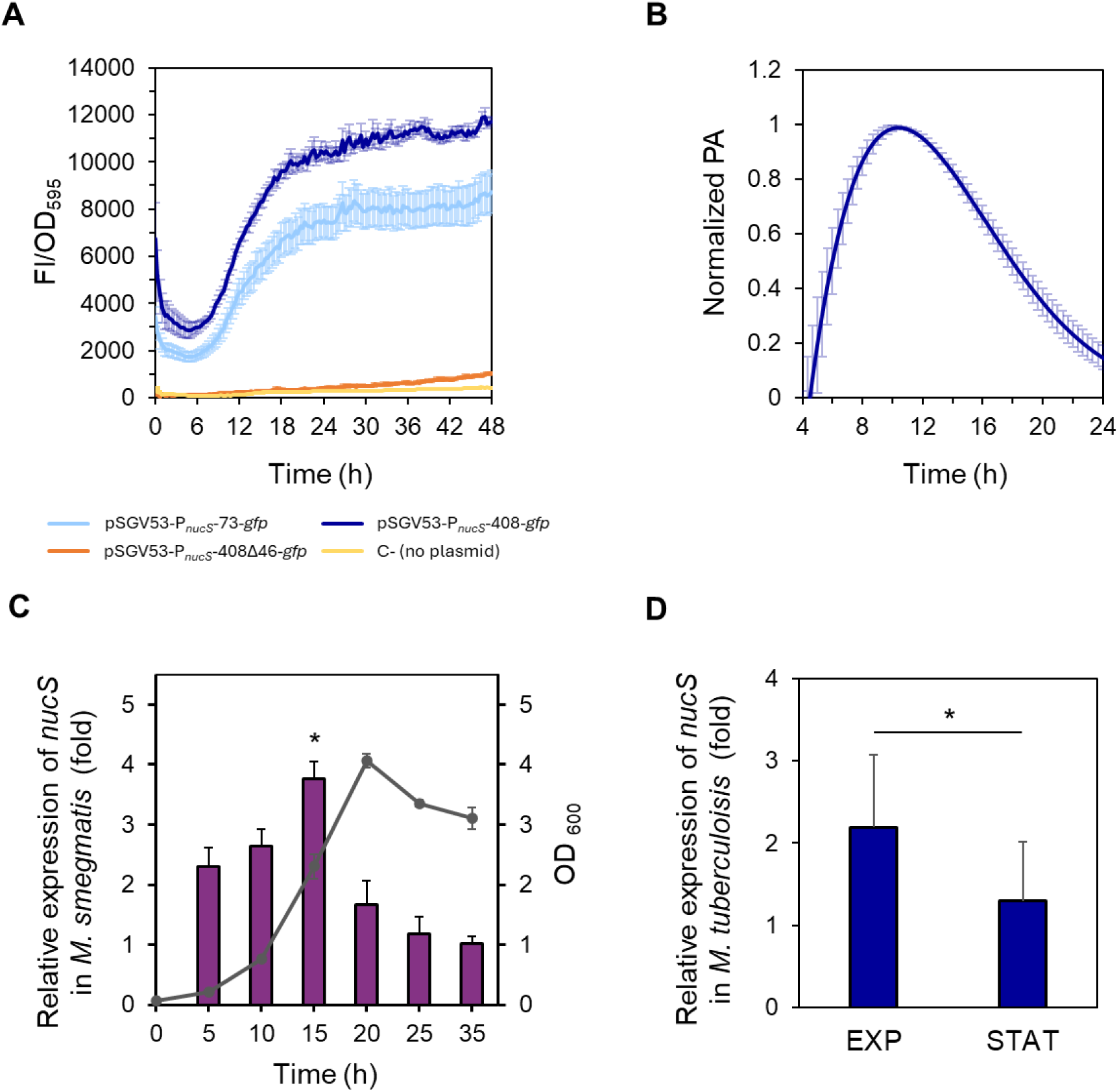
Study of *nucS* expression during growth A. Spectrofluorometric analysis of reporter strains during growth. Normalized fluorescence (FI/OD_595_, arbitrary units) of *M. smegmatis* strains carrying different *nucS::gfp* transcriptional fusions: pSGV53-P*_nucS_*-73-*gfp* (light blue), pSGV53-P*_nucS_*-408-*gfp* (dark blue) and pSGV53-P*_nucS_*-408Δ46-*gfp* (orange). Autofluorescence of non-transformed cells (no plasmid) is shown in yellow (C-, negative control). Data represent the mean ± standard error (SE) of eight biological replicates (n=8). **B. Promoter activity of *M. smegmatis nucS*.** The promoter activity (PA) values of the *M. smegmatis* reporter strain harboring the plasmid pSGV53-P*_nucS_*-408-*gfp* normalized to their respective maximum are shown (see *Materials and Methods*). Error bars: SE (n = 8). **C. Measurement of *nucS* expression in *M. smegmatis* during growth by RT-qPCR.** Bars show the relative expression levels of *nucS* normalized to the endogenous control *sigA* at different growth times. Relative expression was calculated using the 2^-ΔΔCt^ method, with the latest time point (35 h) serving as the control with baseline levels, assigned a value of 1. The growth curve showing OD_600_ values at the different time points when samples were collected is represented in gray. Error bars: SE (n=3). *: p < 0.05 (15 h vs. 35 h), Student’s t-test for paired samples. **D. Measurement of *nucS* expression in *M. tuberculosis* H37Rv at different growth phases.** Bars show the expression levels (2^-ΔΔCt^ method) of *nucS* normalized to the endogenous control *lpqM* at exponential phase (OD_600_=1) relative to the value at stationary phase (STAT) (OD_600_=5). Error bars: SE (n=3). *: p < 0.05, Student’s t-test for paired samples.

When comparing the fluorescence levels of the short (73 bp) and long (408 bp) fusions, we observed that the fluorescence was moderately higher (approximately 30–40%) in the strains carrying the long fusion, suggesting the presence of additional regulatory elements between 74 and 408 bp upstream of the *nucS* TSS. However, the presence of the short region alone was sufficient to drive high fluorescence levels (**Figure 3A**). In contrast, the strain carrying the vector pSGV53-P*_nucS_*-408Δ46-*gfp* exhibited fluorescence levels comparable to the non-transformed negative control (**Figure 3A**), indicating that the intergenic region between *nucS* and the upstream gene is essential for *nucS* expression. These results further support the location of the promoter within this region.

Focusing on the fluorescence pattern during bacterial growth of the reporter strains carrying the short and long fusions, we found that fluorescence intensity steadily increased during the exponential phase in both strains. This suggests that the highest expression of *nucS* occurs during this phase, coinciding with active cell division. Fluorescence continued to increase into the early stationary phase, reaching maximum levels, and then remained constant throughout the rest of the stationary phase (**Figure 3A**). This stable signal is likely due to the long half-life of GFP (23, 24), which allows fluorescence to accumulate even after transcriptional activity has decreased.

To quantify promoter activity (PA), the rate of GFP accumulation was calculated from the spectrofluorometric data (see *Materials and Methods*). Maximum promoter activity for the reporter fusion with the longest upstream region of *nucS* (pSGV53-P*_nucS_*-408-*gfp*) was observed approximately at 10 h of growth under the conditions tested in the microplate reader, coinciding with the late exponential phase (see **Figure S1**). Following this peak, the PA gradually declined (**Figure 3B**).

The assays performed with the reporter fusions suggested that the highest expression of *nucS* occurred during the exponential phase. The transcriptional profile of *nucS* throughout the growth cycle was validated by RT-qPCR, confirming that *nucS* expression is growth-phase-dependent (**Figure 3C**). In this assay, the highest levels of *nucS* transcript were observed towards the end of the exponential phase (15 h vs. 35 h expression ratio of 3.8-fold, p=0.0105) (**Figure 3C**). However, when the cells entered the stationary phase, *nucS* transcription experienced a significant decrease, with the gene expression levels remaining low for the rest of the stationary phase (**Figure 3C**). A similar pattern was observed in *M. tuberculosis*, where *nucS* transcription levels were higher during the exponential phase compared to the stationary phase (p=0.0172) (**Figure 3D**).

It was also analyzed whether the decrease in *nucS* expression observed at the transcript level during the stationary phase was noticeable at the protein level. To investigate this, we performed Western blot analyses using protein extracts from *M. smegmatis* collected at different growth times (**Figure 4A**). To normalize NucS expression levels, the expression of the constitutive protein FtsZ, which remains at similar levels during both the exponential and stationary growth phases (25), was also measured. Similar to the transcript levels, NucS protein abundance was higher in the exponential phase than in the stationary phase, particularly when comparing the end of the exponential phase to the late stationary phase (18 h vs. 36 h expression ratio of 2.3-fold, p=0.0207) (**Figure 4B**).

**Figure 4.**
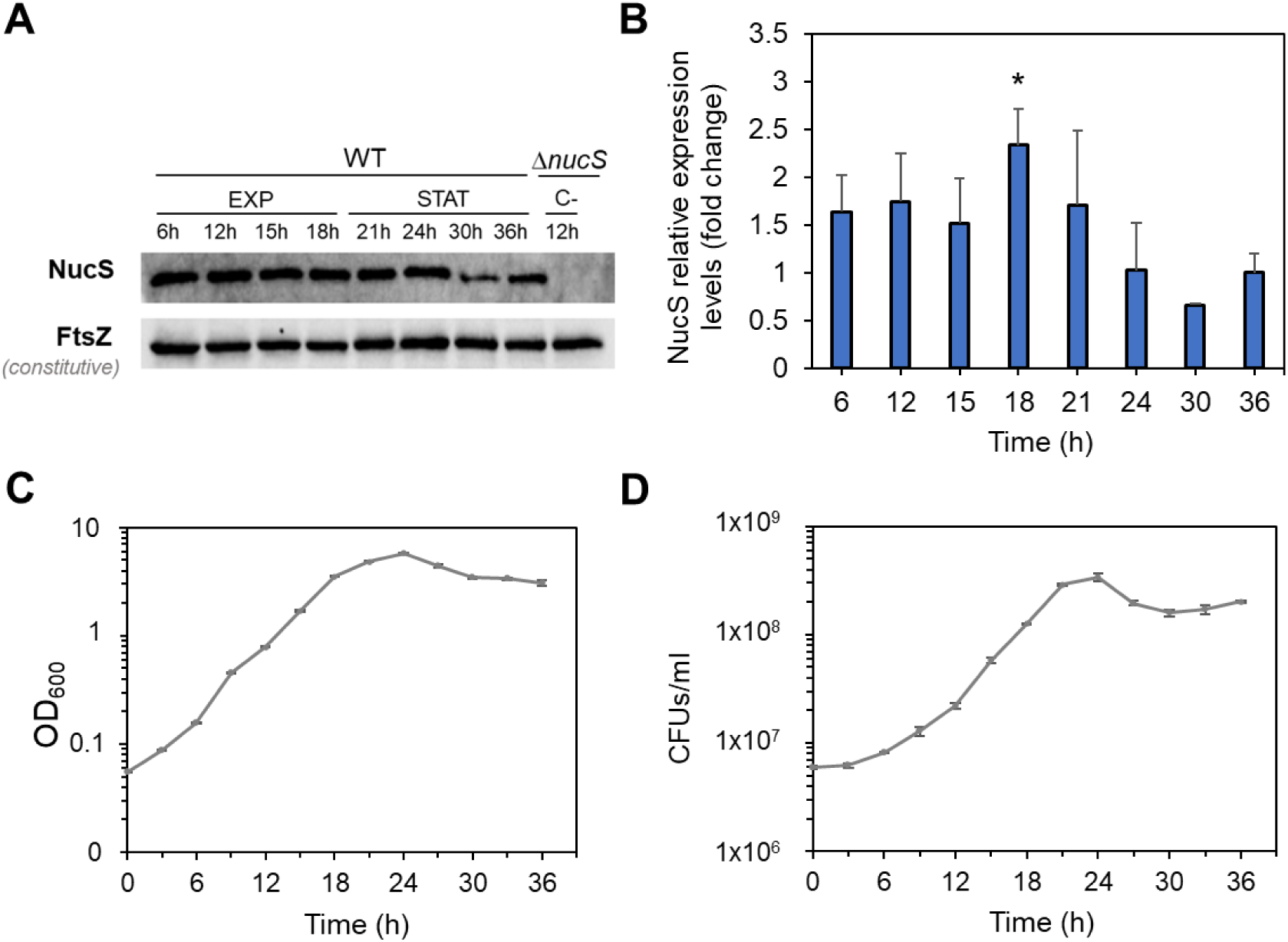
NucS levels in *M. smegmatis* during bacterial growth. A. Western blot analysis of NucS expression. **A.** Representative Western blot experiment showing NucS and FtsZ (loading control) bands in wild-type *M. smegmatis* (WT) at various time points during exponential (EXP) and stationary (STAT) phases. A Δ*nucS* extract collected at 12 h was included as a negative control. Images of the full membrane are provided in **Figure S2**. **B. Quantification of NucS levels.** NucS band intensities were normalized to FtsZ and expressed relative to the 36h time point (baseline=1). Error bars represent SE from three independent experiments using extracts from different biological replicates. *: p < 0.05 (18 h vs. 36 h), paired Student’s t-test. **C. Growth curve of *M. smegmatis* based on optical density.** OD_600_ measurements of the cultures used for the Western blot analysis. Error bars represent SE (n=3). **D. Viability curve based on colony-forming units (CFU).** Average CFU/ml for each time point. Error bars represent SE (n=3).

In summary, the analyses performed using reporter fusions, RT-qPCR and Western blot, indicate that NucS expression at both the transcript and protein levels is growth-phase dependent, decreasing during the stationary phase.

### Analysis of *nucS* expression in a *M. smegmatis sigB*-deficient strain

Reduced expression of canonical MMR system components during the stationary phase has been linked to negative regulation by *rpoS*, which encodes the σ^S^ factor, in microorganisms like *E. coli* (2, 4, 5). In mycobacteria, the gene *sigB*, encoding the sigma factor σ^B^, is functionally analogous to *rpoS* and, like *rpoS*, is induced during the stationary phase and under stress conditions (13–16). Moreover, two putative consensus sequences for σ^B^ binding (NNGNNG) were identified in the upstream region of *nucS* (see **Figure 2B**). This observation suggests that *sigB* may play a role in the regulation of *nucS* during the stationary phase. To investigate this, we compared *nucS* expression in a Δ*sigB* mutant with that of the wild-type strain.

Given our hypothesis that *sigB* negatively regulates *nucS* expression during the stationary phase, we anticipated higher *nucS* expression in the Δ*sigB* mutant compared to the wild-type strain during this phase. RT-qPCR analysis revealed no differences in *nucS* expression between the two strains during the exponential phase. However, during the stationary phase, *nucS* expression in the Δ*sigB* strain was approximately twofold higher than in the wild-type strain (p=0.0259), suggesting negative regulation of *nucS* by *sigB* in the stationary phase (**Figure 5**).

**Figure 5.**
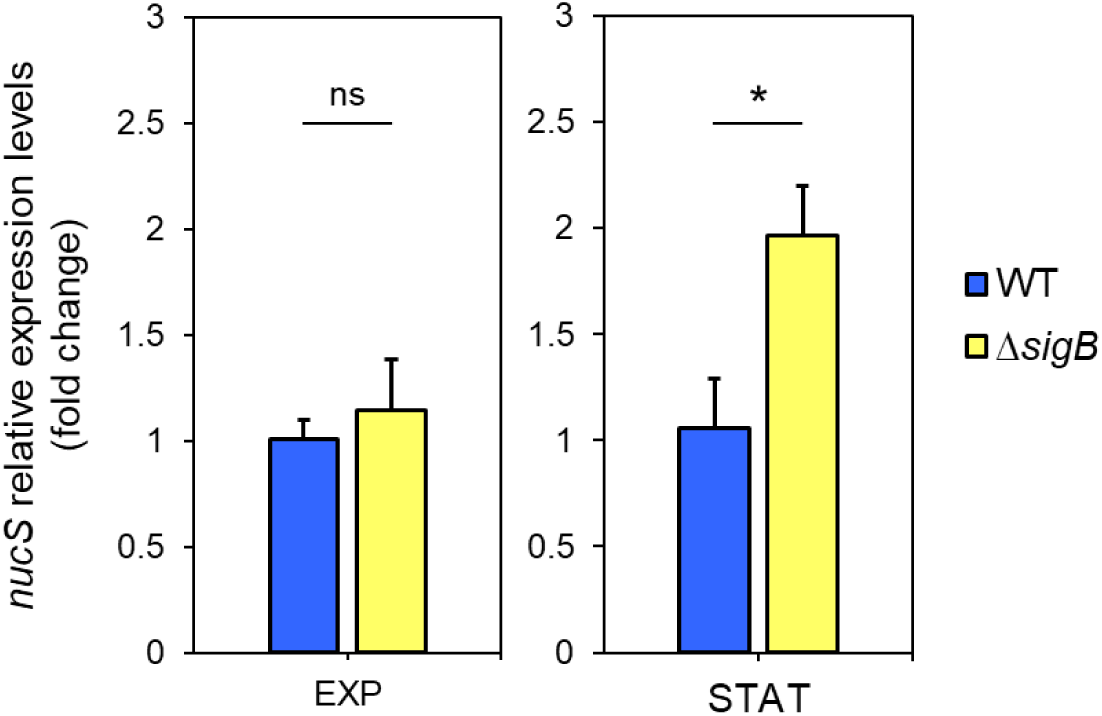
Expression of *nucS* in *M. smegmatis* Δ*sigB* during exponential and stationary phases. Expression levels of *nucS* in the Δ*sigB* mutant relative to the wild-type strain during late exponential (EXP, 15 h) and stationary (STAT, 40 h) phases, as determined by RT-qPCR. Cultures were initiated at OD_600_ = 0.05, and both strains exhibited similar growth rates (see **Figure S3**). Bars represent *nucS* expression normalized to the endogenous control *sigA*. Relative expression was calculated using the 2^-ΔΔCt^ method, with the wild-type strain set as the reference (value = 1). Error bars: SE (n=3). Statistical significance was assessed using a one-tailed t-test for independent samples (alternative hypothesis: Δ*sigB* > wild type; specified as “greater” in R). ns: not significant (p > 0.05); *: p < 0.05.

Additionally, to assess whether the potential regulation of *nucS* by *sigB* during the stationary phase affects the phenotype, the mutant frequency in the presence of rifampicin was calculated for the WT and Δ*sigB* strains during both the exponential and stationary phases. The results showed a higher mutant frequency during the exponential phase compared to the stationary phase for both strains. While the mutant frequency in the WT and Δ*sigB* strains was similar during the exponential phase, the Δ*sigB* strain exhibited a slight (nearly twofold) reduction in mutant frequency relative to the WT strain during the stationary phase, although this difference was not statistically significant (**Figure 6**). Further studies are required to confirm the relevance of potential *nucS* regulation by the σ^B^ factor in the mutator phenotype.

**Figure 6.**
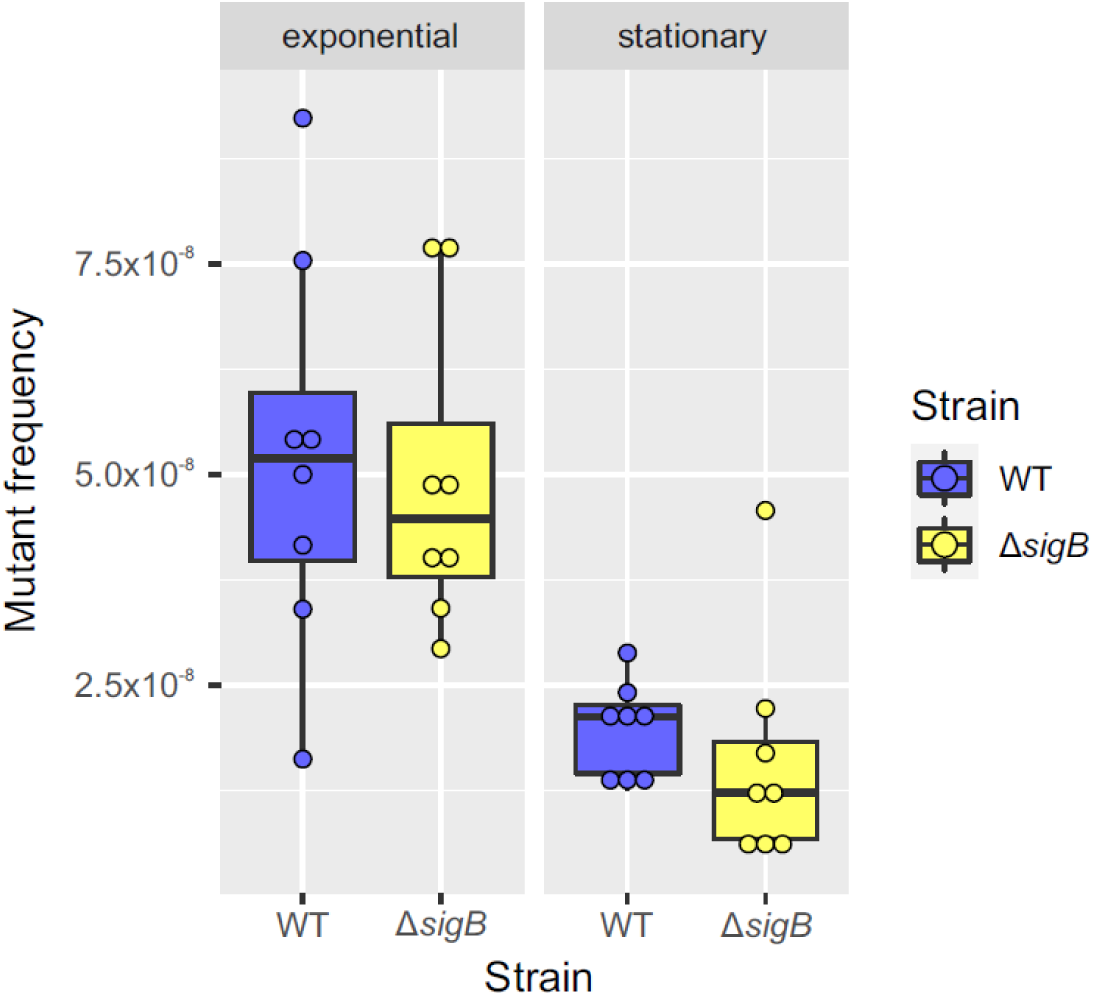
Comparison of mutant frequency between WT and Δ*sigB* strains during exponential and stationary phases. Box plots comparing mutant frequencies between the WT (blue) and Δ*sigB* (yellow) strains during the exponential and stationary phases. The thick black line inside each box represents the median, and the dots show mutant frequency values for each replicate (n=8). No significant differences in mutant frequencies were observed between the two strains in either the exponential or stationary phase (p > 0.05; Mann-Whitney U test). The plots were generated using the ggplot2 package (v3.3.3) in R (v4.0.5).

### Screening for compounds regulating *nucS* expression

Fluorescent reporter fusions are powerful tools for investigating gene expression under diverse conditions. To identify potential compounds regulating *nucS* expression in *M. smegmatis*—either inducers or repressors—a screening was performed using Phenotype MicroArrays (PMs) (Biolog™) in combination with a reporter strain carrying the vector pSGV-P*_nucS_*-408-*gfp* (**Figure 2**, **Table S1**). Biolog™ PMs, designed to evaluate bacterial responses to a wide array of chemical agents, have been successfully applied with fluorescent reporter fusions in prior studies (26). For this screening, chemical sensitivity assay plates (PM11-PM20) were selected. Each plate contained 24 compounds, tested at four increasing concentrations, resulting in the analysis of 240 compounds (**Table S2**).

To establish baseline fluorescence, normalized fluorescence levels (FI/OD_595_) of the reporter strain were measured under screening conditions in the absence of compounds. Upper and lower thresholds were then defined to identify compounds influencing *nucS* expression (**Figure S4**). The upper threshold was calculated as *x* + 0.5*x* + *SD*, where x represents the normalized fluorescence value of the reporter strain at each point, and SD is the standard deviation. The lower threshold was defined as *x* − 0.5*x* − *SD*. Compounds producing fluorescence values above the upper threshold were considered potential inducers, while those yielding values below the lower threshold were classified as potential repressors. This screening identified 22 candidate compounds: 13 putative inducers and 9 putative repressors (**Table S3**).

A follow-up screening was conducted using a disk diffusion assay with the reporter strain and the candidate compounds. This method allows testing a wide range of compound concentrations. Induction of *nucS* expression was observed as an increase in fluorescence intensity around the inhibition halo, while repression was indicated by a decrease in fluorescence in the same region.

Through this assay, only two compounds showed a specific effect on *nucS* expression, producing an increase or decrease in fluorescence relative to the negative control (disk with solvent only) that is not observed in the constitutive expression control (carrying the pSGV53 vector). Specifically, 8-hydroxyquinoline was identified as a potential inducer **(Figure 7A)**, and thioctic acid as a potential repressor **(Figure 7B)**. 8-hydroxyquinoline is a quinoline-family antibiotic with metal-chelating properties (27) and has been reported to exhibit potential mutagenic effects (28). Thioctic acid, also known as α-lipoic acid, also exerts metal-chelating capacity and is primarily recognized for its antioxidant properties, acting as a free radical scavenger and contributing to the repair of oxidative damage (29, 30).

**Figure 7.**
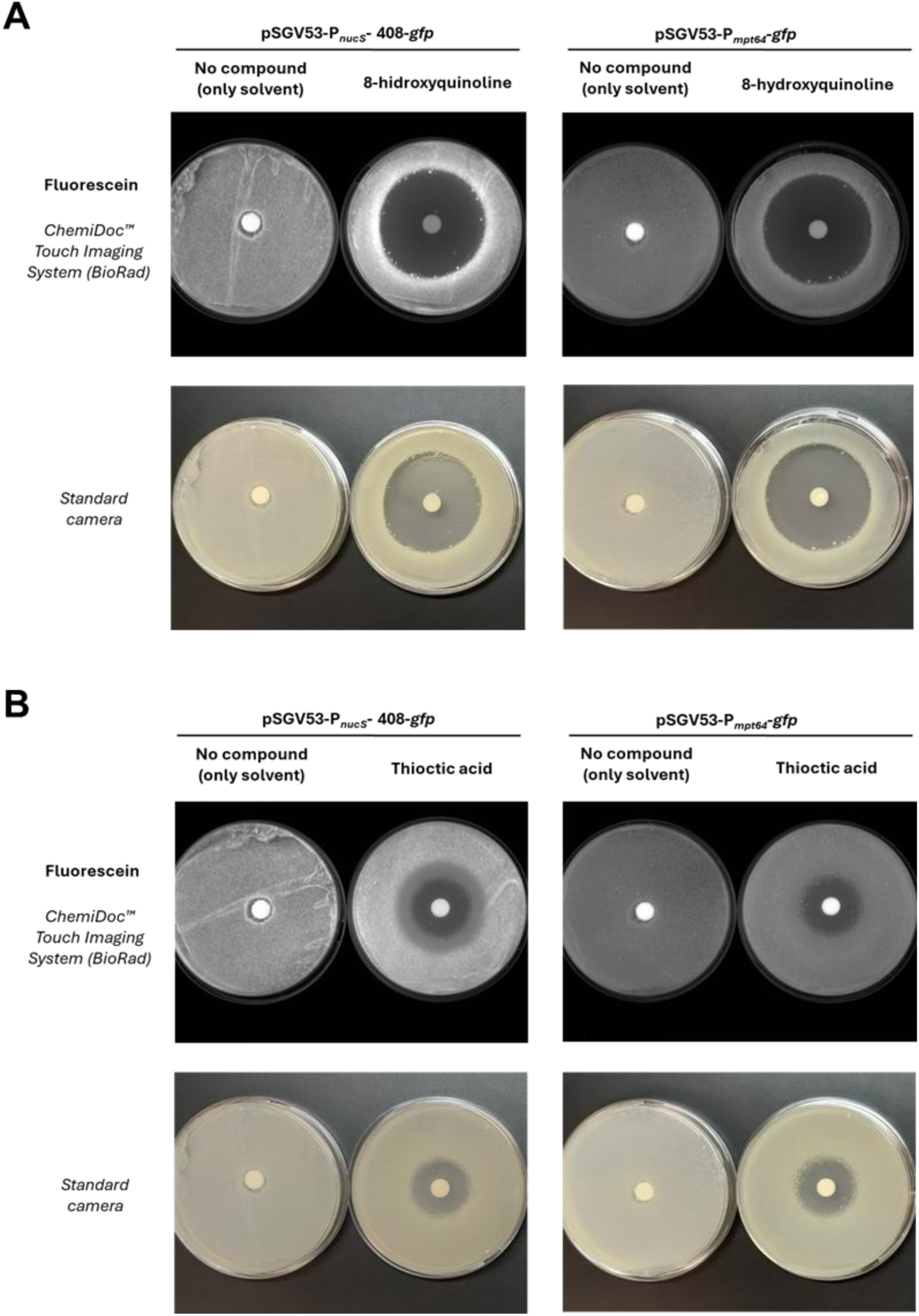
Effect of candidate compounds on *nucS* expression assessed by disk diffusion assay. A. Results with 8-hydroxyquinoline, identified as a potential inducer of *nucS* expression. B. Results with thioctic acid, identified as a potential repressor. The upper pannels (fluorescence images) show the effect of each compound on the expression of the *nucS::gfp* transcriptional fusion in *M. smegmatis*. Induction is observed as increased fluorescence around the inhibition halo, while repression appears as reduced fluorescence in the same region. Disks containing no compound were used as negative controls. A control strain expressing *gfp* under a constitutive promoter (P*_mpt64_*) was included (right side of each pannel) to rule out nonspecific effects. The lower panels display the corresponding photographs of the plates, showing cell mass distribution.

In contrast, other compounds tested produced changes in fluorescence around the inhibition halo in both the reporter and control strains, indicating nonspecific effects of protein expression unrelated to *nucS* regulation **(Figure S5)**.

## DISCUSSION

Variations in the expression of the *nucS* gene may influence the bacterial mutation rate, thereby impacting the acquisition of antibiotic resistance in mycobacterial pathogens, including *M. tuberculosis*. Despite its importance, the regulation and expression of this gene had not been thoroughly studied.

In this study, we identified the transcription start site (TSS) for the *nucS* gene in *M. smegmatis*, which enabled us to identify several candidate –10 regions for sigma factor binding. These included sequences recognized by the primary sigma factor σ^A^ and the alternative stationary phase sigma factor σ^B^ (19). To monitor *nucS* expression, we constructed three reporter strains carrying *nucS*::*gfp* transcriptional fusions with different lengths of upstream regions. Our findings revealed that the minimal 73 bp upstream fragment was sufficient to drive basal *nucS* expression, suggesting that the core promoter is situated near the TSS. However, the higher fluorescence observed with the 408 bp fusion implies that additional regulatory elements between 74 and 408 bp may enhance expression. In contrast, the absence of detectable fluorescence from the fusion lacking the immediate upstream intergenic region (46 bp) underscores the critical role of this segment as the key promoter element. The use of these reporter fusions is an important first step in decoding the regulatory architecture of the *nucS* promoter and exploring regulatory mechanisms governing *nucS* expression. Further studies will be essential to address their specific roles in controlling *nucS* expression.

Our analyses further show that *nucS* expression in *M. smegmatis* is strongly dependent on the growth phase. We observed the highest promoter activity during exponential growth, with a marked reduction in the stationary phase. This finding was corroborated by RT-qPCR and Western blot analyses, all of which suggested that *nucS* transcript and protein level correlate with the physiological state of the cell. During exponential growth, when the rates of cell division and DNA replication are high (31), elevated NucS levels may be required to correct replication errors. Conversely, the reduced replication rate in the stationary phase aligns with lower *nucS* expression. These observations are consistent with previous global transcriptomic studies in both *M. smegmatis* and *M. tuberculosis* (32–34), as well as with the known downregulation of canonical MMR components during periods of reduced cellular replication (2, 3). Altogether, the correlation between NucS levels, growth phase, and replication rates aligns with the behavior expected for an MMR protein that is functionally coupled to DNA replication.

Drawing a parallel with *E. coli*, where the canonical MMR genes *mutS* and *mutH* system are downregulated during the stationary phase via *rpoS* and *hfq* (2, 4, 5), we explored whether the mycobacterial functional analog σ^B^ factor (encoded by *sigB*) (13–16), might similarly influence *nucS* expression (in mycobacteria, no *hfq* homolog has been identified) (35). In a Δ*sigB* mutant of *M. smegmatis*, we observed an approximately twofold increase in *nucS* transcription during stationary phase. Additionally, the Δ*sigB* strain exhibited a roughly two-fold lower mutation frequency than the wild type during the same phase, suggesting that an increase of two-fold in nucS transcription is enough to decrease mutation rate at the same level. These results support the idea that σ^B^ may modulate mutation rates, at least in part, by influencing *nucS* transcription. Whether σ^B^ exerts its effect directly—via binding competition with σ^A^—or indirectly through σ^B^-dependent effectors remains to be determined. It is possible that the downregulation of *nucS* in the stationary phase involves an intermediate regulator whose transcription is promoted by σ^B^, similar to the negative regulation of *mutS* by *rpoS* in *E. coli* during the stationary phase (2). Moreover, the low conservation of mycobacterial promoter sequences, especially in the –35 region, could facilitate the exchange of sigma factors in response to different stress conditions (18, 19, 36). Given the complexity of the mycobacterial transcriptional machinery, with multiple sigma factors (28 in *M. smegmatis* and 13 in *M. tuberculosis*) (14, 19, 37), further experiments are warranted to elucidate these mechanisms at genetic and biochemical level.

Beyond growth phase regulation, our data, along with literature and database analysis (38–42), indicate that *nucS* expression is likely modulated by additional endogenous and exogenous factors. For instance, an antisense ncRNA targeting *nucS* has been reported in *M. tuberculosis* under nitric oxide exposure (12), suggesting a layer of regulation. Similarly, canonical MMR components such as *mutS* have been shown to respond to DNA-damaging agents like mitomycin C (43) and to subinhibitory concentrations of β-lactam antibiotics, which downregulate *mutS* expression in *E. coli* via the induction of the *rpoS* regulon under diverse stress conditions (6). These conditions include nutrient limitation, high osmolarity, extreme temperatures, and low pH (13, 44).

To further identify potential regulators of *nucS* expression, we screened 240 compounds from the Biolog™ bacterial chemical sensitivity plates, using a reporter strain harboring the vector pSGV-P*_nucS_*-408-*gfp*. From this screen, 22 candidate compounds were identified. A subsequent disk diffusion assay highlighted only two compounds with significant effect on *nucS* expression: the quinoline-family antimicrobial 8-hydroxyquinoline, identified as a potential inducer, and the potent antioxidant thioctic acid (α-lipoic acid), which may act as a repressor. 8-hydroxyquinoline is of both natural (plant-derived) and synthetic origin and has been used as a fungicide in agriculture. Thioctic acid, in contrast, is endogenously produced by plants, animals and humans. Both compounds have multiple potential medical applications (27, 29, 30). The precise mechanisms by which these compounds influence *nucS* regulation require further clarification. However, these findings support the hypothesis that *nucS* expression may be modulated by both environmental and/or host-derived factors. Notably, the co-administration of different molecules, as well as patient-specific physiological conditions, may influence bacterial mutation rate and should be considered when designing effective antimicrobial therapies. Beyond the Biolog™ compounds tested here, future studies could explore the effects of additional molecules on *nucS* expression, including DNA-damaging agents, mutagens, antimicrobials, and oxidative stress inducers, using the reporter fusions developed in this study.

In conclusion, our work provides foundational insights into the regulation of *nucS* expression in Mycobacteria, demonstrating its growth-phase dependency and initial responsiveness to potential chemical modulators. These findings underscore the role of NucS as an integral component of the mismatch repair system and suggest that its dynamic regulation may be critical for modulating mutation rates, with possible implications for antibiotic resistance. Additionally, the existence of two distinct pathways at the molecular and evolutionary levels—the *nucS*-mediated non-canonical and the MutS/MutL-mediated canonical MMR systems—which perform almost identical functions, is the result of convergent evolution. Moreover, the fact that these two systems are also regulated in a similar manner provides an example of double homoplasy, both in activity and regulation, across two genetic traits. Future studies should focus on delineating the detailed transcriptional and post-transcriptional mechanisms controlling *nucS* expression, as well as exploring the impact of these regulatory pathways in pathogenic mycobacteria such as *M. tuberculosis*.

## MATERIALS AND METHODS

### Bacterial strains

In this study, we used the *M. smegmatis* mc² 155 (American Type Culture Collection, 700084) wild-type (WT) strain, its noncanonical MMR-deficient (Δ*nucS*) derivative (10) and the Δ*sigB* mutant (this study). *M. tuberculosis* H37Rv was also used in some experiments. *E. coli* DH5α strain was used to obtain recombinant plasmids.

### Culture media and growth conditions

For liquid cultures of *M. smegmatis*, Middlebrook 7H9 broth (Difco) supplemented with 0.5% glycerol and 0.5% Tween 80 was used. All cultures were grown in Erlenmeyer flasks with a 1:5 medium-to-flask volume ratio and incubated at 37°C with orbital shaking (250 rpm). For solid cultures of *M. smegmatis*, Middlebrook 7H10 agar (Difco) supplemented with 0.5% glycerol and 0.05% Tween 80 was used.

Liquid cultures of *M. tuberculosis* were grown in 7H9 broth supplemented with 0.5% glycerol, 0.05% Tween 80, 10% OADC (Oleic Albumin Dextrose Catalase), and incubated in roller bottles at 37°C. The 7H10 agar plates were further supplemented with 5 µg/ml amphotericin.

*E. coli* was cultured in LB broth or LB agar at 37°C.

### Extraction of total RNA from *M. smegmatis*

A volume of 1–2 ml was collected from various *M. smegmatis* cultures under the appropriate conditions for each experiment. After centrifugation (5 min, 12,000 x g), the supernatant was discarded, and the pellets were stored at –80°C for further processing. Initially, cells were washed with 300 μl of TE buffer (10 mM Tris-HCl, 1 mM EDTA, pH 8) and then resuspended in 300–400 μl of lysis buffer. The suspensions were transferred to 2 ml BeadBug™ tubes containing 0.1 mm zirconium beads. Cell lysis was achieved using two cycles of 1 min at maximum speed, with 2 min on ice between cycles, in a BeadBug™ microtube homogenizer. After cell lysis, total RNA was extracted using the RNeasy Mini Kit, Part 1 (QIAGEN), following the manufacturer’s instructions. Residual genomic DNA was removed from the RNA samples using DNase treatment with the Turbo DNA-Free Kit (Invitrogen). RNA concentration was measured with a NanoDrop® (Thermo Fisher Scientific), and sample integrity was confirmed by electrophoresis on a 1.2% agarose gel (100 V, 20– 30 min).

### Extraction of total RNA from *M. tuberculosis*

40-mL cultures were collected at exponential (OD_600_ = 1) and stationary (OD_600_ = 5) phases. Cell pellets were resuspended in 6M guanidine chloride for inactivation and stored at –80°C for a minimum of 7 days. Subsequently, cells were harvested and resuspended in 1 mL of TRIzol™ reagent (Invitrogen), then transferred to 2 ml BeadBug™ tubes containing 0.1 mm zirconium beads. Mechanical cell disruption was performed using a BeadBug™ microtube homogenizer with three cycles of 1 min at 400 rpm, with 2 minutes intervals on ice between cycles. RNA extraction was carried out following the protocol of Rustad *et al.* (45), with minor modifications. Genomic DNA was removed using the DNase I recombinant, RNase-free (Merck), followed by phenol/chloroform extraction to eliminate DNase, and RNA was subsequently precipitated with ethanol.

## 5’-RACE

The transcription start site (TSS) was determined using the 5’-RACE technique, following the protocol by Scotto-Lavino and colleagues (46). Total RNA from *M. smegmatis* RNA was extracted as previously described using the RNeasy Mini Kit, Part 1 (QIAGEN). Reverse transcription was performed using a reverse primer that hybridizes to the internal region of *nucS* (GSP-RT) (**Table S4**). RNAse H (New England Biolabs, NEB) was then added to destroy the RNA template, and the resulting cDNA was purified and polyadenylated at the 3’ end using terminal deoxynucleotidyl transferase (Tdt, NEB) and dATP.

Next, the first round of cDNA amplification was performed by PCR using Q_T_, Q_O_, and GSP_1_ primers (**Table S4**). The PCR product was diluted 1:20 and subjected to a second round of amplification to increase the yield of the specific product using the nested primers QI and GSP2 (**Table S4**). PCR product was visualized by electrophoresis on a 1% agarose gel. The resulting band was purified using the AccuPrep® Gel Purification Kit (BIONEER) and sequenced to determine the TSS.

### Transformation of *M. smegmatis*

Transformation of *M. smegmatis* was performed by electroporation, following the protocol of Goude and Parish (47). A total of 200 ng (for replicative or integrative plasmid) to 5 µg (for homologous recombination) of DNA (in a volume not exceeding 5 µl), previously dialyzed using MF-Millipore® MCE Membrane Filter (0.025 µm) (Merck Millipore), was added to a 200 µl aliquot of electrocompetent cells. The cell-DNA mixture was transferred to pre-chilled 0.2 cm electroporation cuvettes (BioRad) and given a single electric pulse (2.5 kV, 25 µF, 1000 Ω) using a Gene Pulser Xcell™ electroporator (BioRad). After the pulse, the cells were incubated on ice for 10 minutes, then transferred to a flask containing 5 ml of antibiotic-free 7H9 medium and incubated at 37°C for 3 hours. Finally, appropriate volumes (0.1 – 5 ml) were plated on 7H10 agar plates supplemented with the selection antibiotic and incubated at 37°C for 3 to 5 days until colonies appeared.

### Construction of GFP reporter plasmids

Upstream regions of different lengths from the *nucS* gene were amplified by PCR from *M. smegmatis* genomic DNA using the following primer pairs: 73up_nucS_NotI_F/nucS_up_NdeI_R, 408up_nucS_NotI_F/nucS_up_NdeI_R and 408up_nucS_NotI_F/msmeg4924startR (**Table S4**). The amplified fragments corresponded to (i) 73 bp upstream of *nucS* (“short region”), (ii) 408 bp upstream of *nucS* (“long region”), and (iii) a 408 bp region upstream of *nucS* excluding the first 46 bp (which correspond to the *nucS* upstream intergenic region). These fragments were cloned into the replicative plasmid pSGV53 using the restriction enzymes NotI-HF (NEB) and NdeI (NEB) to generate the vectors pSGV53-P*_nucS_*-73-*gfp*, pSGV53-P*_nucS_*-408-*gfp* and pSGV53-P*_nucS_*-408Δ46-*gfp* (**Table S1**). The correct sequences were verified by sequencing using the primers ble_fw_1 and gfp_rev_seq_2 (**Table S4**).

### Spectrofluorometric assays for monitoring fluorescence during growth

Fluorescence and OD_595_ measurements of *M. smegmatis* strains carrying GFP reporter plasmids were measured during growth using an Infinite® 200 spectrofluorometer (TECAN). Black, clear-bottom 96-well plates (Corning™) were used, inoculating 8 wells (replicates) with each strain, with an initial OD_595_ of 0.05 – 0.1 (measured in the spectrofluorometer) in a final volume of 200 μl of 7H9 medium per well. The i-control™ software version 1.6 (TECAN) was used to take measurements every 20 minutes over a 48-hour growth period (145 cycles) at 37°C. The program for each cycle was as follows: i) orbital shaking for 10 s (5 mm amplitude), ii) a waiting time of 15 s, iii) absorbance measurement at 595 nm (25 flashes), iv) fluorescence intensity (FI) measurement: excitation wavelength 485 nm, emission wavelength 530 nm, 25 flashes, Bottom mode, integration time 20 μs, manual gain of 100. Relative fluorescence (FI/ OD_595_) was plotted by dividing the fluorescence intensity by the OD_595_ at each point.

Promoter activity (PA) was calculated following the method described by Camas *et al.* (48). Briefly, relative fluorescence (FI/OD_595_) data were fitted to a sixth-order polynomial, and the derivative of the resulting curves was obtained. PA values were then normalized to their respective maximum.

### Measurement of *nucS* expression by RT-qPCR

For *M. smegmatis*, samples of 1–2 ml were taken from three independent cultures (biological replicates) of each strain under the specified conditions and time points for each case. Total RNA was extracted as previously described. The reverse transcription (RT) reaction was performed using 250 ng of RNA per sample in a final volume of 20 μl with the High-Capacity cDNA Reverse Transcriptase Kit (Applied Biosystems). Quantitative PCR (qPCR) was carried out using Power SYBR® Green PCR Master Mix (Applied Biosystems) on a 7500 Real-Time PCR System (Applied Biosystems). Reactions were prepared in MicroAmp Optical 96-Well Reaction Plates (Applied Biosystems), with each containing 20 μl of Master Mix (10 μl of Power SYBR® Green, 9.2 μl of water, and 0.4 μl of each primer at 25 μM) and 5 μl of cDNA (5 ng/μl). Three biological and three technical replicates were used. The thermocycler conditions were: i) 2 min at 50°C, ii) 10 min at 95°C, iii) 40 cycles of 15 s at 95°C, followed by 1 min at 66°C. Results were analyzed using the 2^−ΔΔCT^ method (121), with the *sigA* housekeeping gene used for normalization.

For *M. tuberculosis*, total RNA was extracted from three independent cultures at exponential (OD_600_ = 1) or stationary phase (OD_600_ = 5), as previously described. Reverse transcription was performed using 5 μg of RNA in a final volume of 80 μl with the High-Capacity cDNA Reverse Transcriptase Kit (Applied Biosystems). qPCR reactions included 5 μl of a 1:25 cDNA dilution (2.5 ng/μl) and 20 μl of Master Mix (12.5 μl of Power SYBR® Green, 7.1 μl of nuclease-free water, and 0.2 μl of each primer at 10 μM). Three biological and three technical replicates were used per condition. Reactions were run on a 7500 Real-Time PCR System (Applied Biosystems) under the following conditions: i) 2 min at 50°C, ii) 10 min at 95°C, iii) 40 cycles of 15 s at 95°C, followed by 1 min at 60°C. Relative expression was calculated using the 2^−ΔΔCT^ method (121), with *lpqM* gene used as the reference gene, as it has been shown to maintain constant expression levels across many different conditions in *M. tuberculosis* (49). The sequences of the primers used in the qPCR reactions are listed in **Table S4**.

### Generation of a polyclonal mouse anti-NucS antibody

For antigen production, the first 294 bp of the *nucS* gene from *M. smegmatis* were cloned into the pET-24a(+) vector (Novagen) using the restriction enzymes NdeI and XhoI. The resulting recombinant 12 kDa N-terminal fragment of NucS was expressed in *E. coli* BL21(DE3) and purified by affinity chromatography using TALON® Metal Affinity Resin (TaKaRa). The purified protein fragment was subsequently coupled to KLH following standard procedures. A polyclonal mouse antibody against NucS from *M. smegmatis* was generated by immunizing female BALB/c mice (n = 5/group; 6-8 weeks-old at the beginning of the study) with three subcutaneous doses (25 µg/dose) of the KLH-conjugated antigen. On day 14 after the third antibody boost, blood was collected from each mouse by submandibular bleeding to obtain serum samples that were stored at –20°C until analysis of humoral immune responses. A pool of anti-NucS antibody was generated by mixing equal amounts of serum from mice selected for their high antibody titer. The mouse studies were approved by the Ethical Committee of Animal Experimentation (CEEA) of the CNB-CSIC, the CSIC Ethics Committee and the Division of Animal Protection of the Comunidad de Madrid (PROEX 184.0_21). Animal procedures conformed with international guidelines and with Spanish law under the Royal Decree (RD) 53/2013. The mouse studies were conducted in accordance with the local legislation and institutional requirements.

### Generation of a polyclonal anti-FtsZ antibody

For antigen production, the *ftsZ* gene from *M. tuberculosis* was cloned into plasmid pET-28a(+) (Novagen) using the restriction enzymes NcoI and XhoI. Polyclonal antibodies against mycobacterial FtsZ, capable of detecting FtsZ from both *M. tuberculosis* and *M. smegmatis*, were generated in rabbits by Charles River Laboratories using their standard immunization protocol.

### Analysis of NucS protein levels in *M. smegmatis* by Western Blot

Samples of 1.5–12 ml were taken from three independent cultures of *M. smegmatis* at the indicated time points and centrifugated to pellet the cells, which were stored at –80°C until further processing. For cell lysis, pellets were resuspended in ∼300 μl of PBS supplemented with cOmplete™ protease inhibitor cocktail (Roche) (25X) and sonicated (minimum of 4 cycles of 20 pulses at 0.7 s). Protein concentration was measured using the Pierce™ BCA Protein Assay Kit (ThermoFisher Scientific).

Proteins were separated by SDS-PAGE using the Mini-PROTEAN III vertical electrophoresis system (BioRad) on 12% resolving gels and 4% stacking gels. A total of 20 μg of protein per sample was mixed with loading buffer (4X) and 0.35 M DTT, heated at 100°C for 10 min, and separated by electrophoresis in Tris-Glycine-SDS running buffer (25 mM Tris, 190 mM glycine, 0.1% SDS) at 200 V for 5 minutes, then at 130 V for about 1 hour and 20 minutes. Transfer to an Immobilon-P PVDF membrane (0.45 μm) (Merck Millipore) was performed using the Trans-Blot® SD Semi-Dry Transfer Cell (BioRad) for 70 min at 15 V.

The membrane was blocked by incubation with gentle agitation in PBS – 0.05% Tween 20 for 30 minutes, followed by PBS – 0.05% Tween 20 – 1% BSA for 1 hour, and finally in PBS – 0.05% Tween 20 – 1% BSA – 5% milk powder for 30 minutes. The membrane was then incubated overnight at 4°C with the polyclonal mouse anti-NucS antibody (1/8000). After washing in PBS – 0.05% Tween 20, the membrane was incubated for 1 hour with goat anti-mouse (IgG-h+I, HRP conjugated) secondary antibody (1:2500, Bethyl Laboratories) at room temperature. After washing with PBS-0.05% Tween 20 and water, signal detection was performed using ECL Western detection reagents (GE Healthcare) and imaged using a ChemiDoc™ Touch Imaging System (BioRad).

For FtsZ detection (loading control), the membrane was re-incubated with PBS – 0.02% sodium azide to inhibit the HRP peroxidase of the previous secondary antibody and blocked again with PBS – 0.05% Tween 20 – 1% BSA – 5% milk powder. Rabbit anti-FtsZ antibody (1:1500) was incubated overnight at 4°C, followed by detection with HRP-Protein A (1:10000, Invitrogen) for 40 min at room temperature. Band intensity was quantified using ImageJ software (Bio-Formats plugin), and NucS signal was normalized to FtsZ.

### **Growth Curve (OD**_600_ and CFUs/ml) of *M. smegmatis*

Three independent *M. smegmatis* cultures were inoculated at an initial OD_600_ of 0.05 in 500 ml flasks containing 100 ml of medium and incubated at 37°C with shaking at 250 rpm. OD_600_ measurements were taken every 3 hours (approximately the generation time of *M. smegmatis*) over a period of 36 hours using a spectrophotometer, and samples were collected for CFU/ml counts. Appropriate dilutions from each culture were plated in duplicate on 7H10 solid medium to obtain colony counts between 30 and 200 (from 200 x 10^-4^ μl at t=0 to 500 x 10^-6^ μl at t=36 h). Plates were incubated at 37°C for 3–4 days before counting the colonies.

### Construction of the *M. smegmatis* Δ*sigB* mutant

The mutant was constructed following the method of Parish and Stoker (50). Two ∼ 1 kb fragments flanking the *sigB* gene, both upstream (sigB5’) and downstream (sigB3’) of the gene, were amplified by PCR from *M. smegmatis* genomic DNA using the primers sigB5_PstIF/sigB5_HindIIIR and sigB3_HindIIIF/sigB3_BamHIR (**Table S4**). Both fragments were sequentially cloned into the p2NIL plasmid (50) (**Table S1**) using the appropriate restriction enzymes. The final suicide plasmid was generated by inserting the pGOAL19 marker cassette into the unique *Pac*I site of the p2NIL plasmid containing the *sigB* flanking fragments.

*M. smegmatis* was transformed by electroporation with 5 μl of the suicide plasmid (500–1000 ng/μl). Merodiploid selection (single recombination event) was performed on 7H10 medium with kanamycin (25 μg/ml) and X-gal (100 μg/ml). Blue colonies resistant to hygromycin (50 μg/ml) were cultured on 7H10 without antibiotics to promote the second recombination event. After one day of growth, the cultures were transferred to liquid 7H9 medium without antibiotics for another 24 hours. Serial dilutions (10^-1^, 10^-2^, and 10^-3^) were plated on 7H10 with X-gal (100 μg/ml) and 10% sucrose. White colonies (with both WT and Δ*sigB* genotypes) were tested for kanamycin (25 μg/ml) and hygromycin (50 μg/ml) sensitivity. PCR verification of the *sigB* deletion was performed using the primers sigB5_end_F/sigB3_start_R and sigB.intF/sigB.intR (**Table S4**, **Figure S6**).

### Estimation of mutant frequency in different growth phases of *M. smegmatis* wild-type and Δ*sigB* strains

Eight independent cultures per strain were inoculated and incubated overnight. A 1:10000 dilution was performed to avoid the selection of pre-existing rifampicin-resistant mutants. Viable and mutant counts were conducted during the exponential phase (29-32 hours of incubation, OD_600_ = 0.6-0.8) and the stationary phase (52-54 hours OD_600_ = 4-6).

In the exponential phase, 300 μl of a 10^-5^ dilution of all cultures were plated on solid 7H10 medium without antibiotics for viable counts. For mutant counts, 30 ml of culture were plated on 7H10 medium with rifampicin (100 μg/ml).

In the stationary phase, 300 μl of a 10^-6^ dilution were plated on 7H10 without antibiotics for viable counts. For mutant counts, 10 ml of culture were plated on 7H10 medium with rifampicin (100 μg/ml). Plates without antibiotics were incubated for 4 days, and rifampicin plates for 7 days, after which colonies were counted. Mutant frequency per ml for each strain was determined by dividing the number of mutants/ml by the number of viable cells/ml for each culture, and the median was calculated across all cultures.

### Screening for compounds regulating *nucS* expression Using Biolog™ Phenotype MicroArrays

Biolog™ PM11-PM20 plates were used for the compound screening (**Table S2**). To prepare the plates, the lyophilized compounds at the bottom of each well were dissolved in 80 μl of liquid 7H9 medium, followed by gentle shaking for 2–3 hours. The plates were then inoculated with the wild-type *M. smegmatis* strain harboring the plasmid pSGV53-P*_nucS_*-408-*gfp* at an initial OD_595_ of approximately 0.1 (measured using an Infinite® 200 spectrofluorometer [TECAN]) in a final volume of 100 μl of medium per well.

The plates were incubated at 37°C with periodic shaking for 72 hours in an Infinite® 200 spectrofluorometer (TECAN), with measurements of OD_595_ and fluorescence intensity (FI, excitation/emission: 485/530 nm) taken every 10 minutes (433 cycles). Relative fluorescence was calculated as the ratio of fluorescence intensity to OD_595_ over the entire growth period.

To establish fluorescence thresholds for identifying compounds that could induce or repress *nucS* expression, a control assay was performed using the same *M. smegmatis* reporter strain grown under identical conditions in a plate without compounds. The thresholds served as benchmarks to identify wells with fluorescence values indicative of potential overexpression or repression of *nucS*.

### Effect of candidate compounds in *nucS* expression using disk diffusion assays

The effect of candidate compounds from the Biolog™ PM screening on nucS expression was qualitatively evaluated using the *M. smegmatis* strain carrying the reporter plasmid pSGV53-P*_nucS_*-408-*gfp*. To rule out nonspecific effects on protein expression, a control strain harboring the pSGV53 vector (with the *gfp* gene under the constitutive promoter P*_mpt64_*, referred to as pSGV53-P*_mpt64_*-*gfp* in **Figure 7** and **Figure S5**) was included.

Overnight cultures were adjusted to the same OD_600_ and diluted 1:10 before being plated on 7H10 agar using the flood inoculation method. Filter disks (Whatman® Antibiotic Assay Discs diam. 9 mm, Merck) were placed on the agar surface and spotted with 30 μl of compounds stock solutions, prepared with the appropriate solvent and concentration (**Table S3**). Disks containing only the corresponding solvent were used as negative controls. After incubation at 37 °C for 72 h, fluorescence was visualized using a ChemiDoc™ Touch Imaging System (BioRad) under fluorescein settings (Blot > Fluorescein). Photographs of the plates were also taken to provide a visual reference for cell mass distribution.

## ACKNOWLEDGMENTS

This research was supported by grants PCI2019-103488/AEI/10.13039/501100011033 and PID2020-112865RBI00/AEI/10.13039/501100011033 from the State Research Agency (AEI), and FIS PI17/00159 from the Carlos III Institute of Health (ISCIII) and the European Regional Development Fund (FEDER, EU). E.C.-S. is the recipient of a PFIS predoctoral fellowship (FI18/00036), co-funded by the ISCIII and the European Social Fund. A.R.-E. is the recipient of an FPU fellowship from the Ministry of Science, Innovation and Universities (FPU2020/01676).

We thank José R. Valverde and A. Couce for statistical and evolutionary advice, respectively, Renata Moreno for her helpful input on some of the experiments and the staff of the CNB Protein Tools Unit for their help with mouse immunizations and immune response analyses.

## SUPPLEMENTAL MATERIAL

### Supplementary tables

**Table S1.**
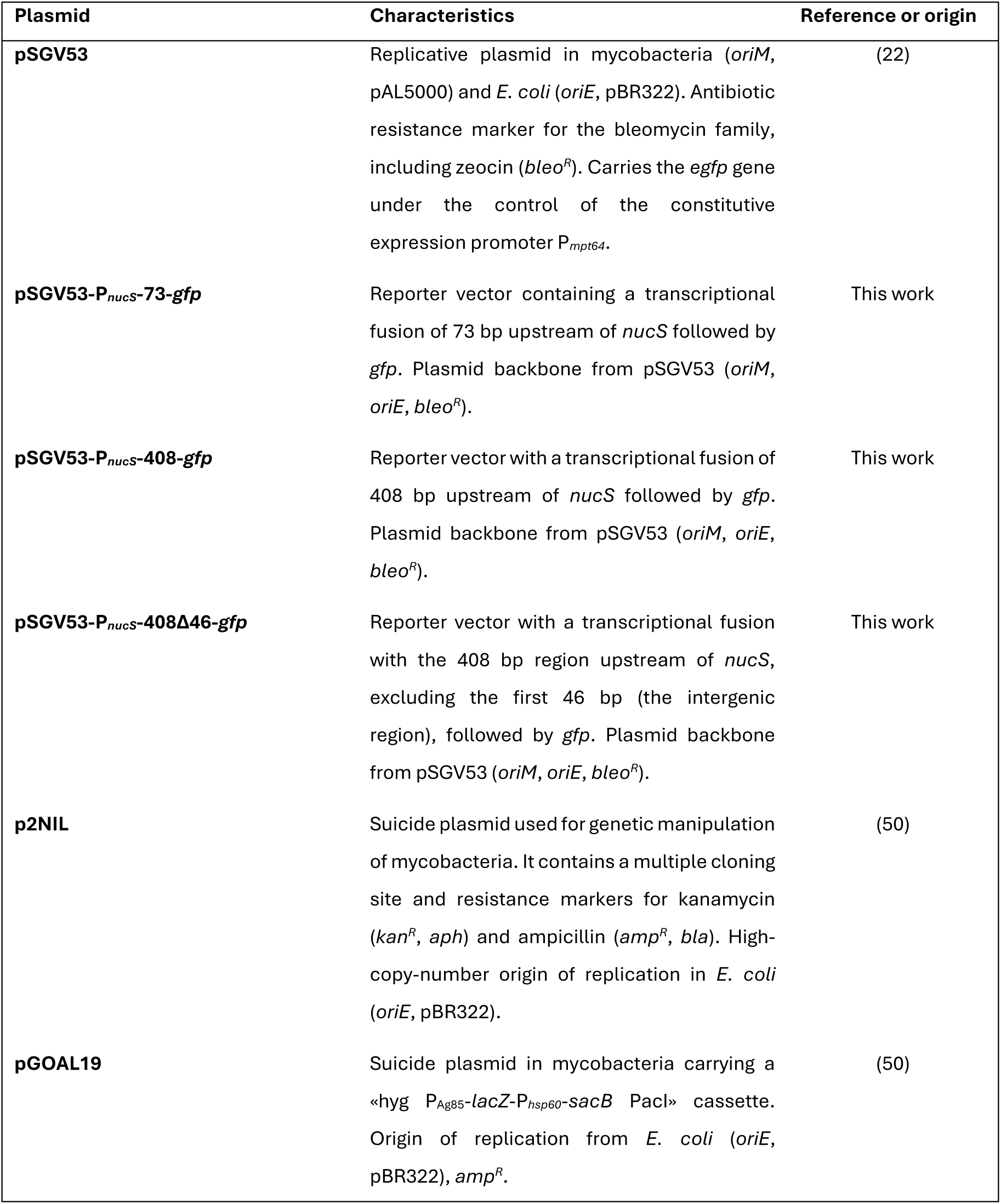
Vectors used in this study.

**Table S2.** Compounds tested from the Biolog™ Phenotype MicroArrays™.

| **Compound** | **Biolog™ Plate** |
| --- | --- |
| Amikacin | *PM11C MicroPlate™* |
| Chlortetracycline | *PM11C MicroPlate™* |
| Lincomycin | *PM11C MicroPlate™* |
| Amoxicillin | *PM11C MicroPlate™* |
| Cloxacillin | *PM11C MicroPlate™* |
| Lomefloxacin | *PM11C MicroPlate™* |
| Bleomycin | *PM11C MicroPlate™* |
| Colistin | *PM11C MicroPlate™* |
| Minocycline | *PM11C MicroPlate™* |
| Capreomycin | *PM11C MicroPlate™* |
| Demeclocycline | *PM11C MicroPlate™* |
| Nafcillin | *PM11C MicroPlate™* |
| Cefazolin | *PM11C MicroPlate™* |
| Enoxacin | *PM11C MicroPlate™* |
| Nalidixic acid | *PM11C MicroPlate™* |
| Chloramphenicol | *PM11C MicroPlate™* |
| Erythromycin | *PM11C MicroPlate™* |
| Neomycin | *PM11C MicroPlate™* |
| Ceftriaxone | *PM11C MicroPlate™* |
| Gentamicin | *PM11C MicroPlate™* |
| Potassium tellurite | *PM11C MicroPlate™* |
| Cephalothin | *PM11C MicroPlate™* |
| Kanamycin | *PM11C MicroPlate™* |
| Ofloxacin | *PM11C MicroPlate™* |
| Penicillin G | *PM12B MicroPlate™* |
| Tetracycline | *PM12B MicroPlate™* |
| Carbenicillin | *PM12B MicroPlate™* |
| Oxacillin | *PM12B MicroPlate™* |
| Penimepicycline | *PM12B MicroPlate™* |
| Polymyxin B | *PM12B MicroPlate™* |
| Paromomycin | *PM12B MicroPlate™* |
| Vancomycin | *PM12B MicroPlate™* |
| D,L-Serine hydroxamate | *PM12B MicroPlate™* |
| Sisomicin | *PM12B MicroPlate™* |
| Sulfamethazine | *PM12B MicroPlate™* |
| Novobiocin | *PM12B MicroPlate™* |
| 2,4-Diamino-6,7-diisopropylpteridine | *PM12B MicroPlate™* |
| Sulfadiazine | *PM12B MicroPlate™* |
| Benzethonium chloride | *PM12B MicroPlate™* |
| Tobramycin | *PM12B MicroPlate™* |
| Sulfathiazole | *PM12B MicroPlate™* |
| 5-Fluoroorotic acid | *PM12B MicroPlate™* |
| Spectinomycin | *PM12B MicroPlate™* |
| Sulfamethoxazole | *PM12B MicroPlate™* |
| L-Aspartic-β-hydroxamate | *PM12B MicroPlate™* |
| Spiramycin | *PM12B MicroPlate™* |
| Rifampicin | *PM12B MicroPlate™* |
| Dodecyltrimethyl ammonium bromide | *PM12B MicroPlate™* |
| Ampicillin | *PM13B MicroPlate™* |
| Dequalinium chloride | *PM13B MicroPlate™* |
| Nickel chloride | *PM13B MicroPlate™* |
| Azlocillin | *PM13B MicroPlate™* |
| 2, 2'-Dipyridyl | *PM13B MicroPlate™* |
| Oxolinic acid | *PM13B MicroPlate™* |
| 6-Mercaptopurine | *PM13B MicroPlate™* |
| Doxycycline | *PM13B MicroPlate™* |
| Potassium chromate | *PM13B MicroPlate™* |
| Cefuroxime | *PM13B MicroPlate™* |
| 5-Fluorouracil | *PM13B MicroPlate™* |
| Rolitetracycline | *PM13B MicroPlate™* |
| Cytosine-1-beta-D-arabinofuranoside | *PM13B MicroPlate™* |
| Geneticin (G418) | *PM13B MicroPlate™* |
| Ruthenium red | *PM13B MicroPlate™* |
| Cesium chloride | *PM13B MicroPlate™* |
| Glycine | *PM13B MicroPlate™* |
| Thallium (I) acetate | *PM13B MicroPlate™* |
| Cobalt chloride | *PM13B MicroPlate™* |
| Manganese chloride | *PM13B MicroPlate™* |
| Trifluoperazine | *PM13B MicroPlate™* |
| Cupric chloride | *PM13B MicroPlate™* |
| Moxalactam | *PM13B MicroPlate™* |
| Tylosin | *PM13B MicroPlate™* |
| Acriflavine | *PM14A MicroPlate™* |
| Furaltadone | *PM14A MicroPlate™* |
| Sanguinarine | *PM14A MicroPlate™* |
| 9-Aminoacridine | *PM14A MicroPlate™* |
| Fusaric acid | *PM14A MicroPlate™* |
| Sodium arsenate | *PM14A MicroPlate™* |
| Boric Acid | *PM14A MicroPlate™* |
| 1-Hydroxypyridine -2-thione | *PM14A MicroPlate™* |
| Sodium cyanate | *PM14A MicroPlate™* |
| Cadmium chloride | *PM14A MicroPlate™* |
| Iodoacetate | *PM14A MicroPlate™* |
| Sodium dichromate | *PM14A MicroPlate™* |
| Cefoxitin | *PM14A MicroPlate™* |
| Nitrofurantoin | *PM14A MicroPlate™* |
| Sodium metaborate | *PM14A MicroPlate™* |
| Chloramphenicol | *PM14A MicroPlate™* |
| Piperacillin | *PM14A MicroPlate™* |
| Sodium metavanadate | *PM14A MicroPlate™* |
| Chelerythrine | *PM14A MicroPlate™* |
| Carbenicillin | *PM14A MicroPlate™* |
| Sodium nitrite | *PM14A MicroPlate™* |
| EGTA | *PM14A MicroPlate™* |
| Promethazine | *PM14A MicroPlate™* |
| Sodium orthovanadate | *PM14A MicroPlate™* |
| Procaine | *PM15B MicroPlate™* |
| Guanidine hydrochloride | *PM15B MicroPlate™* |
| Cefmetazole | *PM15B MicroPlate™* |
| D-Cycloserine | *PM15B MicroPlate™* |
| EDTA | *PM15B MicroPlate™* |
| 5,7-Dichloro- 8-hydroxyquinaldine | *PM15B MicroPlate™* |
| 5,7-Dichloro-8-hydroxyquinoline | *PM15B MicroPlate™* |
| Fusidic acid | *PM15B MicroPlate™* |
| 1,10-Phenanthroline | *PM15B MicroPlate™* |
| Phleomycin | *PM15B MicroPlate™* |
| Domiphen bromide | *PM15B MicroPlate™* |
| Nordihydroguaia retic acid | *PM15B MicroPlate™* |
| Alexidine | *PM15B MicroPlate™* |
| 5-Nitro-2-furaldehyde semicarbazone | *PM15B MicroPlate™* |
| Methyl viologen | *PM15B MicroPlate™* |
| 3, 4-Dimethoxybenzyl alcohol | *PM15B MicroPlate™* |
| Oleandomycin | *PM15B MicroPlate™* |
| Puromycin | *PM15B MicroPlate™* |
| CCCP | *PM15B MicroPlate™* |
| Sodium azide | *PM15B MicroPlate™* |
| Menadione | *PM15B MicroPlate™* |
| 2-Nitroimidazole | *PM15B MicroPlate™* |
| Hydroxyurea | *PM15B MicroPlate™* |
| Zinc chloride | *PM15B MicroPlate™* |
| Cefotaxime | *PM16A MicroPlate™* |
| Phosphomycin | *PM16A MicroPlate™* |
| 5-Chloro-7-iodo-8-hydroxyquinoline | *PM16A MicroPlate™* |
| Norfloxacin | *PM16A MicroPlate™* |
| Sulfanilamide | *PM16A MicroPlate™* |
| Trimethoprim | *PM16A MicroPlate™* |
| Dichlofluanid | *PM16A MicroPlate™* |
| Protamine sulfate | *PM16A MicroPlate™* |
| Cetylpyridinium chloride | *PM16A MicroPlate™* |
| 1-Chloro -2,4-dinitrobenzene | *PM16A MicroPlate™* |
| Diamide | *PM16A MicroPlate™* |
| Cinoxacin | *PM16A MicroPlate™* |
| Streptomycin | *PM16A MicroPlate™* |
| 5-Azacytidine | *PM16A MicroPlate™* |
| Rifamycin SV | *PM16A MicroPlate™* |
| Potassium tellurite | *PM16A MicroPlate™* |
| Sodium selenite | *PM16A MicroPlate™* |
| Aluminum sulfate | *PM16A MicroPlate™* |
| Chromium chloride | *PM16A MicroPlate™* |
| Ferric chloride | *PM16A MicroPlate™* |
| L-Glutamic-ghydroxamate | *PM16A MicroPlate™* |
| Glycine hydroxamate | *PM16A MicroPlate™* |
| Chloroxylenol | *PM16A MicroPlate™* |
| Sorbic acid | *PM16A MicroPlate™* |
| D-Serine | *PM17A MicroPlate™* |
| β-Chloro-L-alanine hydrochloride | *PM17A MicroPlate™* |
| Thiosalicylic acid | *PM17A MicroPlate™* |
| Sodium salicylate | *PM17A MicroPlate™* |
| Hygromycin B | *PM17A MicroPlate™* |
| Ethionamide | *PM17A MicroPlate™* |
| 4-Aminopyridine | *PM17A MicroPlate™* |
| Sulfachloropyridazine | *PM17A MicroPlate™* |
| Sulfamonomethoxine | *PM17A MicroPlate™* |
| Oxycarboxin | *PM17A MicroPlate™* |
| 3-Amino-1,2,4-triazole | *PM17A MicroPlate™* |
| Chlorpromazine | *PM17A MicroPlate™* |
| Niaproof | *PM17A MicroPlate™* |
| Compound 48/80 | *PM17A MicroPlate™* |
| Sodium tungstate | *PM17A MicroPlate™* |
| Lithium chloride | *PM17A MicroPlate™* |
| DL-Methionine hydroxamate | *PM17A MicroPlate™* |
| Tannic acid | *PM17A MicroPlate™* |
| Chlorambucil | *PM17A MicroPlate™* |
| Cefamandole nafate | *PM17A MicroPlate™* |
| Cefoperazone | *PM17A MicroPlate™* |
| Cefsulodin | *PM17A MicroPlate™* |
| Caffeine | *PM17A MicroPlate™* |
| Phenylarsine oxide | *PM17A MicroPlate™* |
| Ketoprofen | *PM18C MicroPlate™* |
| Sodium pyrophosphate decahydrate | *PM18C MicroPlate™* |
| Thiamphenicol | *PM18C MicroPlate™* |
| Trifluorothymidin | *PM18C MicroPlate™* |
| Pipemidic Acid | *PM18C MicroPlate™* |
| Azathioprine | *PM18C MicroPlate™* |
| Poly-L-lysine | *PM18C MicroPlate™* |
| Sulfisoxazole | *PM18C MicroPlate™* |
| Pentachlorophenol | *PM18C MicroPlate™* |
| Sodium m-arsenite | *PM18C MicroPlate™* |
| Sodium bromate | *PM18C MicroPlate™* |
| Lidocaine | *PM18C MicroPlate™* |
| Sodium metasilicate | *PM18C MicroPlate™* |
| Sodium m-periodate | *PM18C MicroPlate™* |
| Antimony (III) chloride | *PM18C MicroPlate™* |
| Semicarbazide | *PM18C MicroPlate™* |
| Tinidazole | *PM18C MicroPlate™* |
| Aztreonam | *PM18C MicroPlate™* |
| Triclosan | *PM18C MicroPlate™* |
| 3,5-Diamino-1,2,4-triazole (Guanazole) | *PM18C MicroPlate™* |
| Myricetin | *PM18C MicroPlate™* |
| 5-fluoro-5'- deoxyuridine | *PM18C MicroPlate™* |
| 2-Phenylphenol | *PM18C MicroPlate™* |
| Plumbagin | *PM18C MicroPlate™* |
| Josamycin | *PM19 MicroPlate™* |
| Gallic acid | *PM19 MicroPlate™* |
| Coumarin | *PM19 MicroPlate™* |
| Methyltrioctylammonium chloride | *PM19 MicroPlate™* |
| Harmane | *PM19 MicroPlate™* |
| 2,4-Dinitrophenol | *PM19 MicroPlate™* |
| Chlorhexidine | *PM19 MicroPlate™* |
| Umbelliferone | *PM19 MicroPlate™* |
| Cinnamic acid | *PM19 MicroPlate™* |
| Disulphiram | *PM19 MicroPlate™* |
| Iodonitro Tetrazolium Violet | *PM19 MicroPlate™* |
| Phenyl- methylsulfonylfluoride (PMSF) | *PM19 MicroPlate™* |
| FCCP | *PM19 MicroPlate™* |
| D,L-Thioctic Acid | *PM19 MicroPlate™* |
| Lawsone | *PM19 MicroPlate™* |
| Phenethicillin | *PM19 MicroPlate™* |
| Blasticidin S | *PM19 MicroPlate™* |
| Sodium caprylate | *PM19 MicroPlate™* |
| Lauryl sulfobetaine | *PM19 MicroPlate™* |
| Dihydrostreptomycin | *PM19 MicroPlate™* |
| Hydroxylamine | *PM19 MicroPlate™* |
| Hexamine cobalt (III) chloride | *PM19 MicroPlate™* |
| Thioglycerol | *PM19 MicroPlate™* |
| Polymyxin B | *PM19 MicroPlate™* |
| Amitriptyline | *PM20B MicroPlate™* |
| Apramycin | *PM20B MicroPlate™* |
| Benserazide | *PM20B MicroPlate™* |
| Orphenadrine | *PM20B MicroPlate™* |
| D,L-Propranolol | *PM20B MicroPlate™* |
| Tetrazolium violet | *PM20B MicroPlate™* |
| Thioridazine | *PM20B MicroPlate™* |
| Atropine | *PM20B MicroPlate™* |
| Ornidazole | *PM20B MicroPlate™* |
| Proflavine | *PM20B MicroPlate™* |
| Ciprofloxacin | *PM20B MicroPlate™* |
| 18-Crown-6-ether | *PM20B MicroPlate™* |
| Crystal violet | *PM20B MicroPlate™* |
| Dodine | *PM20B MicroPlate™* |
| Hexachlorophene | *PM20B MicroPlate™* |
| 4-Hydroxycoumarin | *PM20B MicroPlate™* |
| Oxytetracycline | *PM20B MicroPlate™* |
| Pridinol | *PM20B MicroPlate™* |
| Captan | *PM20B MicroPlate™* |
| 3,5-Dinitrobenzene | *PM20B MicroPlate™* |
| 8-Hydroxyquinoline | *PM20B MicroPlate™* |
| Patulin | *PM20B MicroPlate™* |
| Tolylfluanid | *PM20B MicroPlate™* |
| Troleandomycin | *PM20B MicroPlate™* |

**Table S3.**
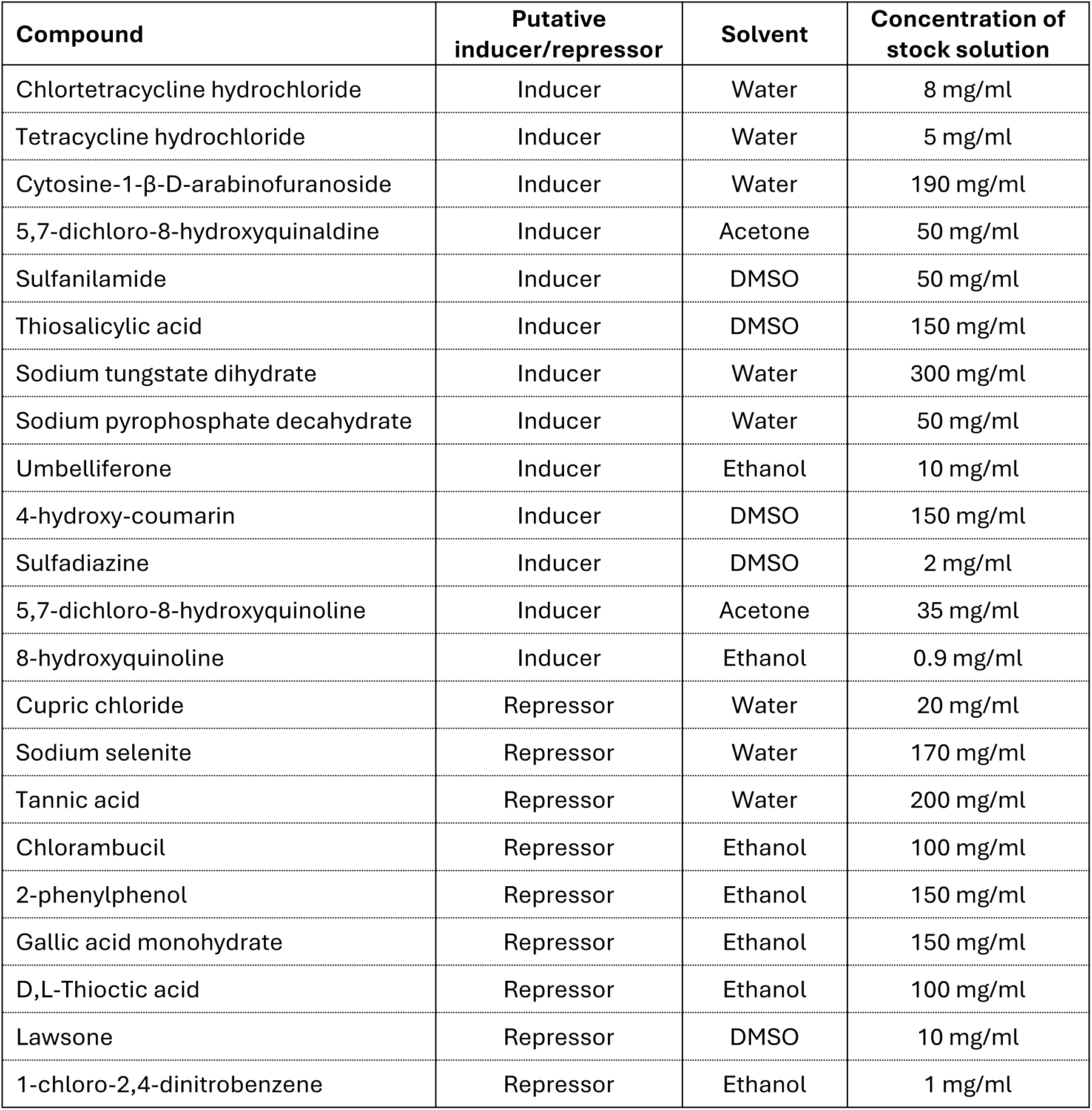
Compounds tested in the disk diffusion assay.

**Table S4.**
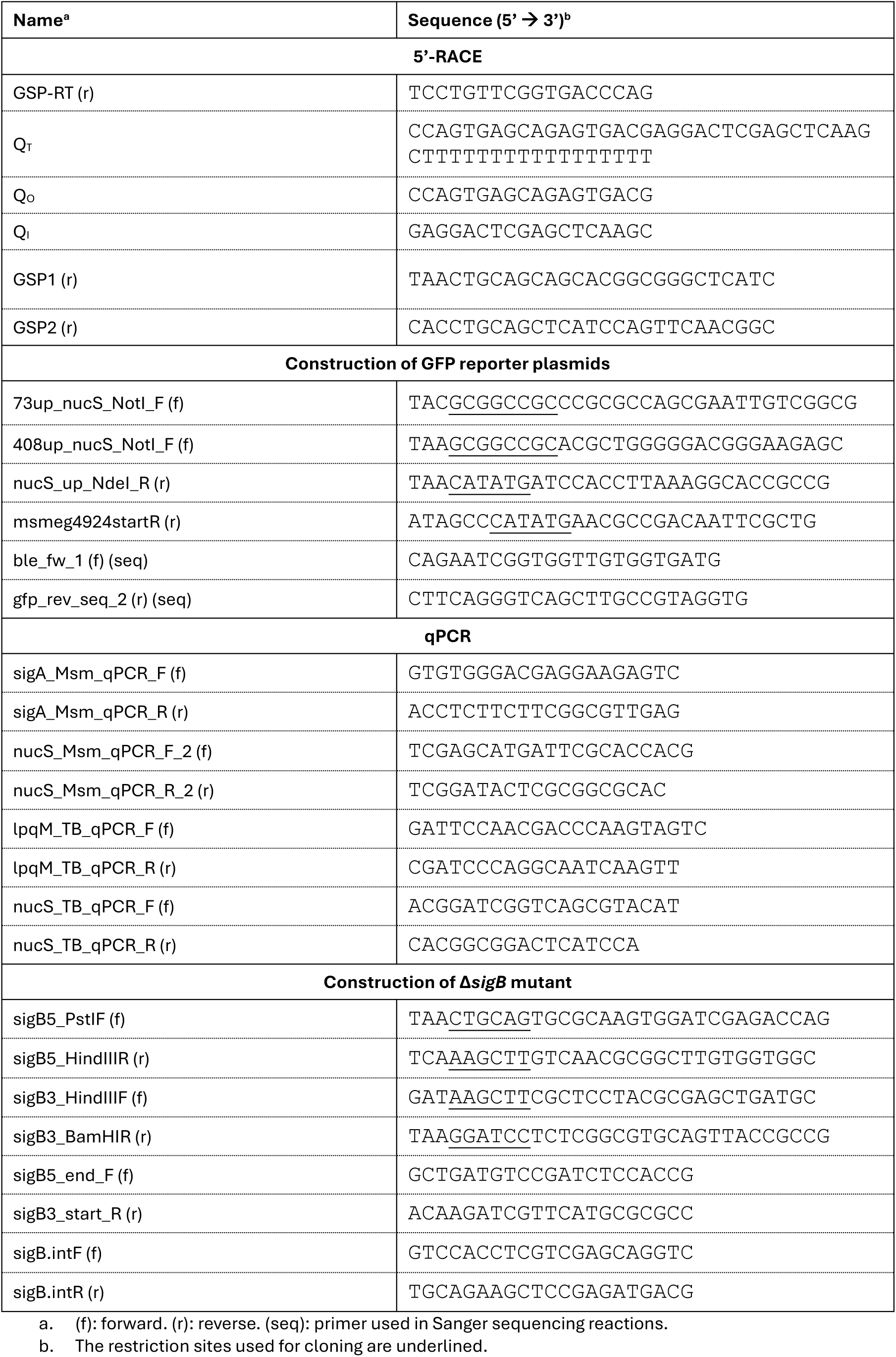
Primers used in this study.

### Supplementary figures

**Figure S1.**
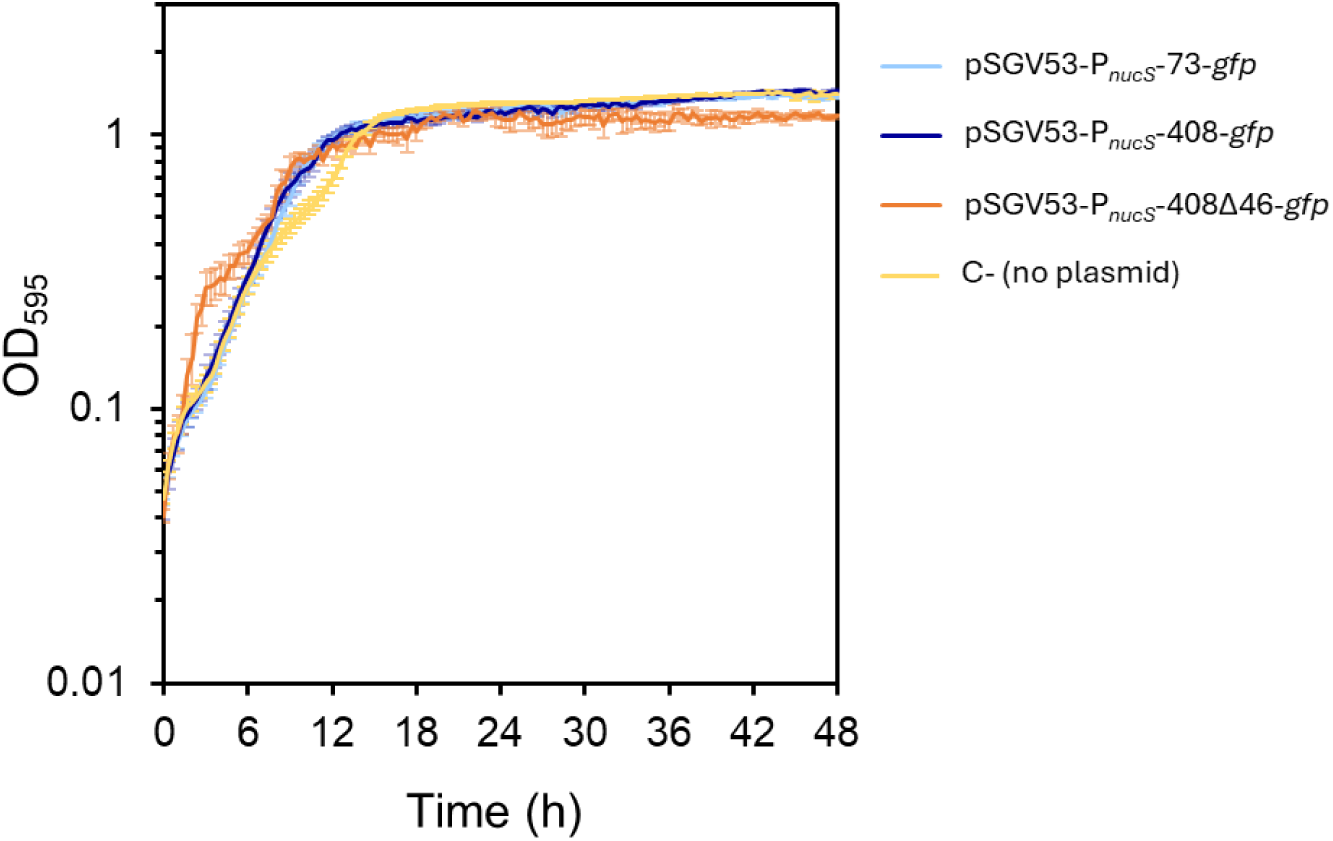
Growth curves of the reporter strains in the microplate reader. OD_595_ values were recorded during growth using an Infinite® 200 spectrofluorometer (TECAN) (see *Materials and Methods*). Data represent the mean ± standard error (SE) of eight biological replicates (n=8).

**Figure S2.**
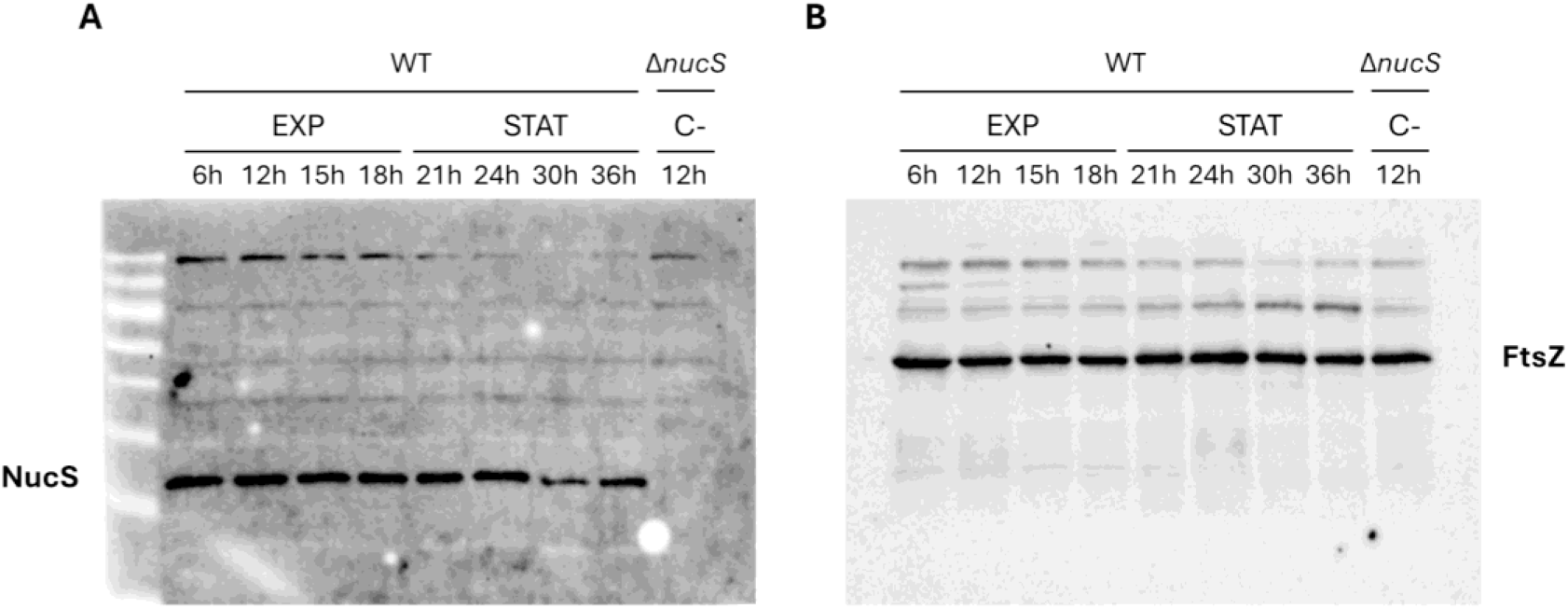
Full membrane images from a representative Western blot experiment. A. Detection of NucS in *M. smegmatis* protein extracts collected at the indicated time points. The membrane was incubated with a mouse anti-NucS antibody followed by a goat anti-mouse IgG-HRP secondary antibody (see *Materials and Methods* section). **B. Detection of FtsZ bands in the same samples, used as a loading control.** After NucS detection, the membrane was treated with sodium azide to inactivate residual peroxidase activity and then incubated with a rabbit anti-FtsZ antibody followed by HRP-Protein A (see *Materials and Methods* section). No NucS signal was observed when HRP-Protein A was used. A Δ*nucS* extract collected at 12 h was included as a negative control. EXP: exponential phase; STAT: stationary phase. Signal detection was performed using ECL reagents and visualized with a ChemiDoc™ Touch Imaging System (Bio-Rad).

**Figure S3.**
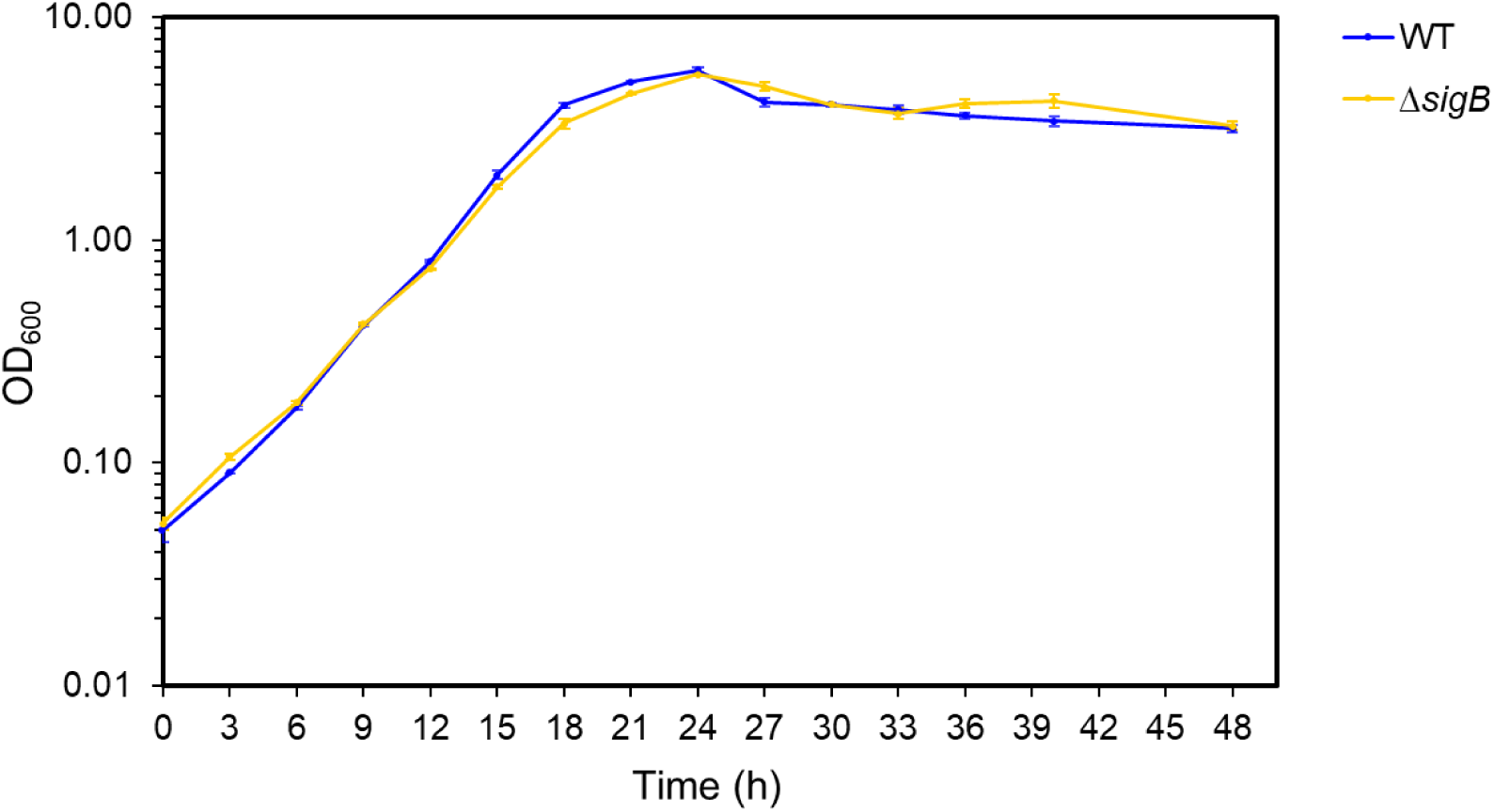
Growth curves of WT and Δ*sigB* strains of *M. smegmatis* mc^2^ 155. Growth of WT (blue) and Δ*sigB* (yellow) strains over 48 h. Both strains exhibited similar growth rates. Cultures were initiated at OD_600_ = 0.05 and incubated at 37°C with orbital shaking (250 rpm). Error bars represent the standard error of three biological replicates.

**Figure S4.**
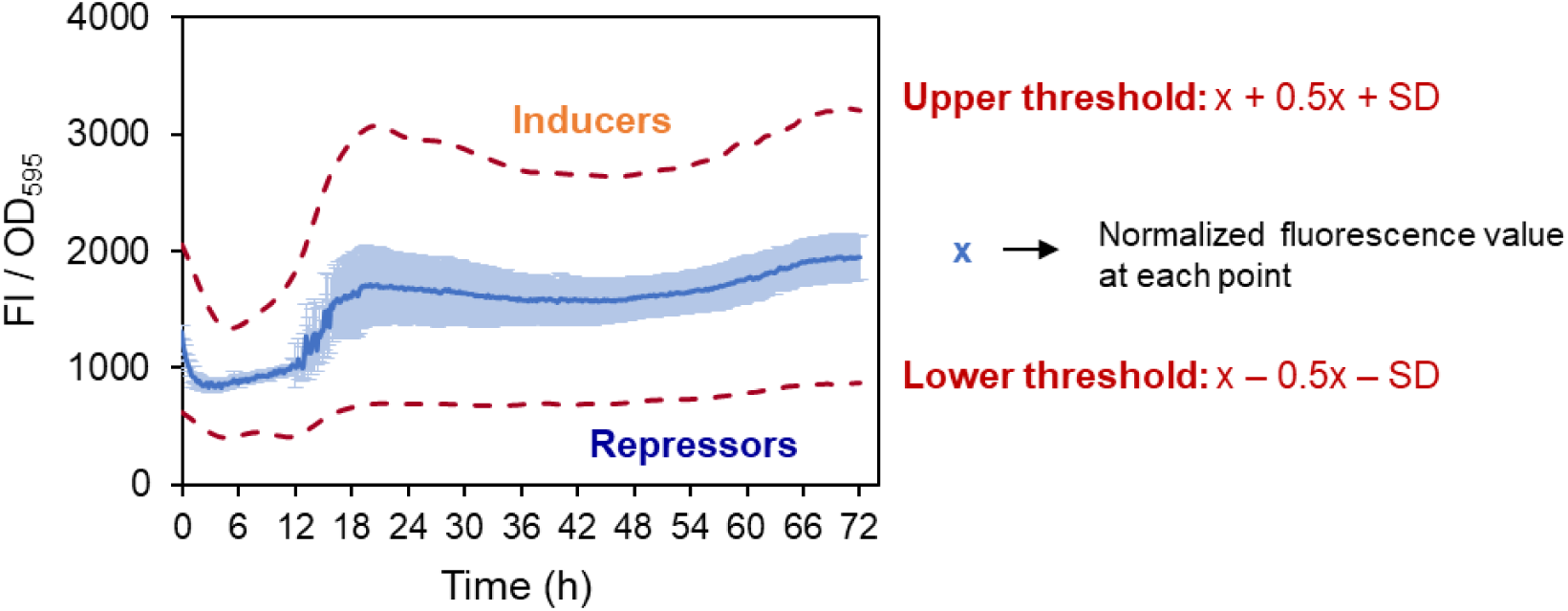
Selection of thresholds for filtering candidate compounds regulating *nucS* expression. The blue line represents the mean normalized fluorescence relative to OD (FI/OD_595_) over 72 hours of growth. Light blue error bars indicate the standard deviation (SD) (n=7). Dashed red lines show the upper and lower threshold values.

**Figure S5.**
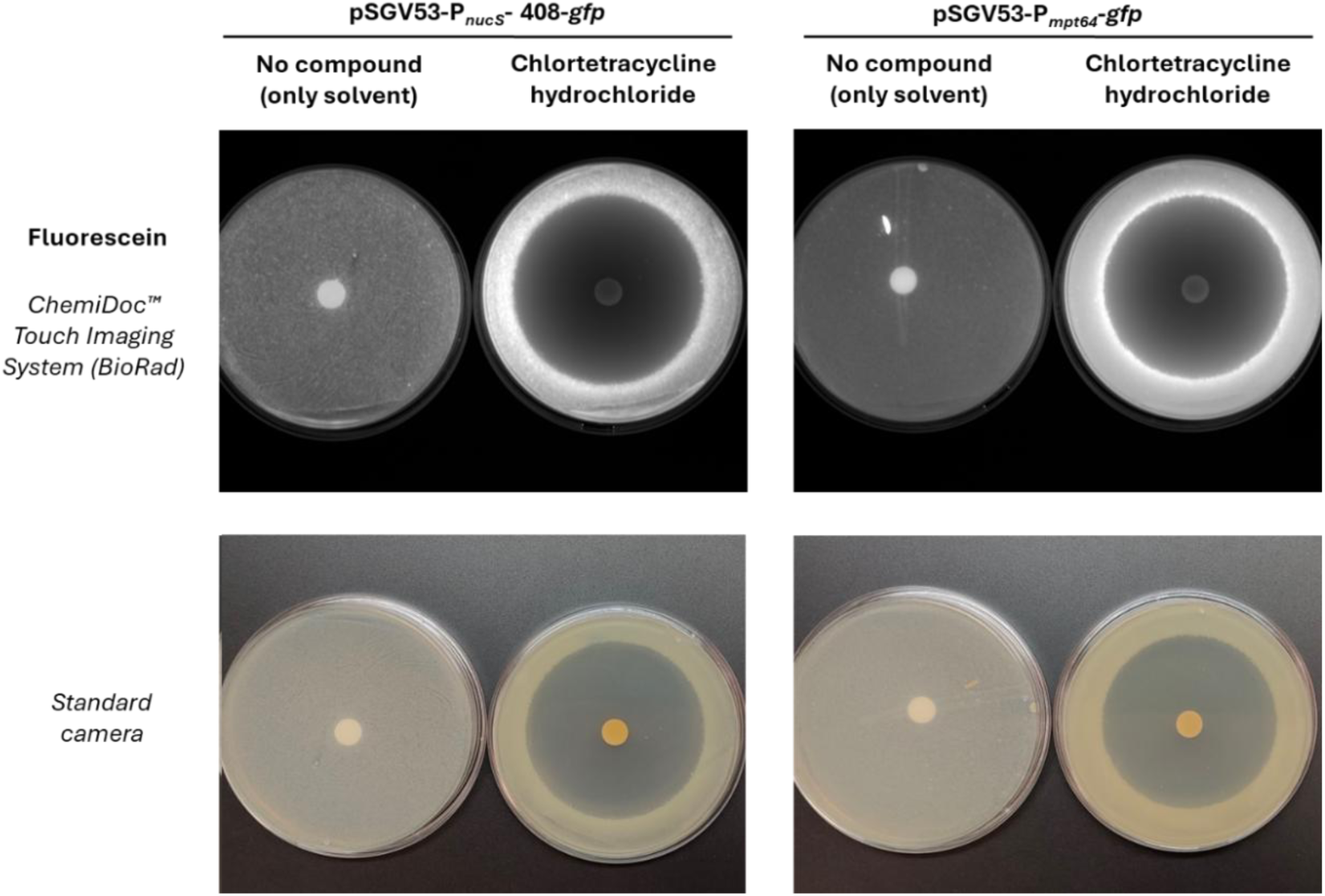
Disk diffusion assay with chlortetracycline, a discarded candidate compound showing nonspecific effects on *nucS* expression. Top panels (fluorescence): GFP fluorescence of the *M. smegmatis nucS::gfp* reporter strain (left panel) and a constitutive GFP-expressing control strain (right panel), each exposed to the test compound (right disk) and solvent control (left disk). Increased fluorescence around the inhibition halo is visible in both strains, indicating a nonspecific effect on protein expression. Bottom panels: Corresponding photographs of the same plates showing cell mass distribution.

**Figure S6.**
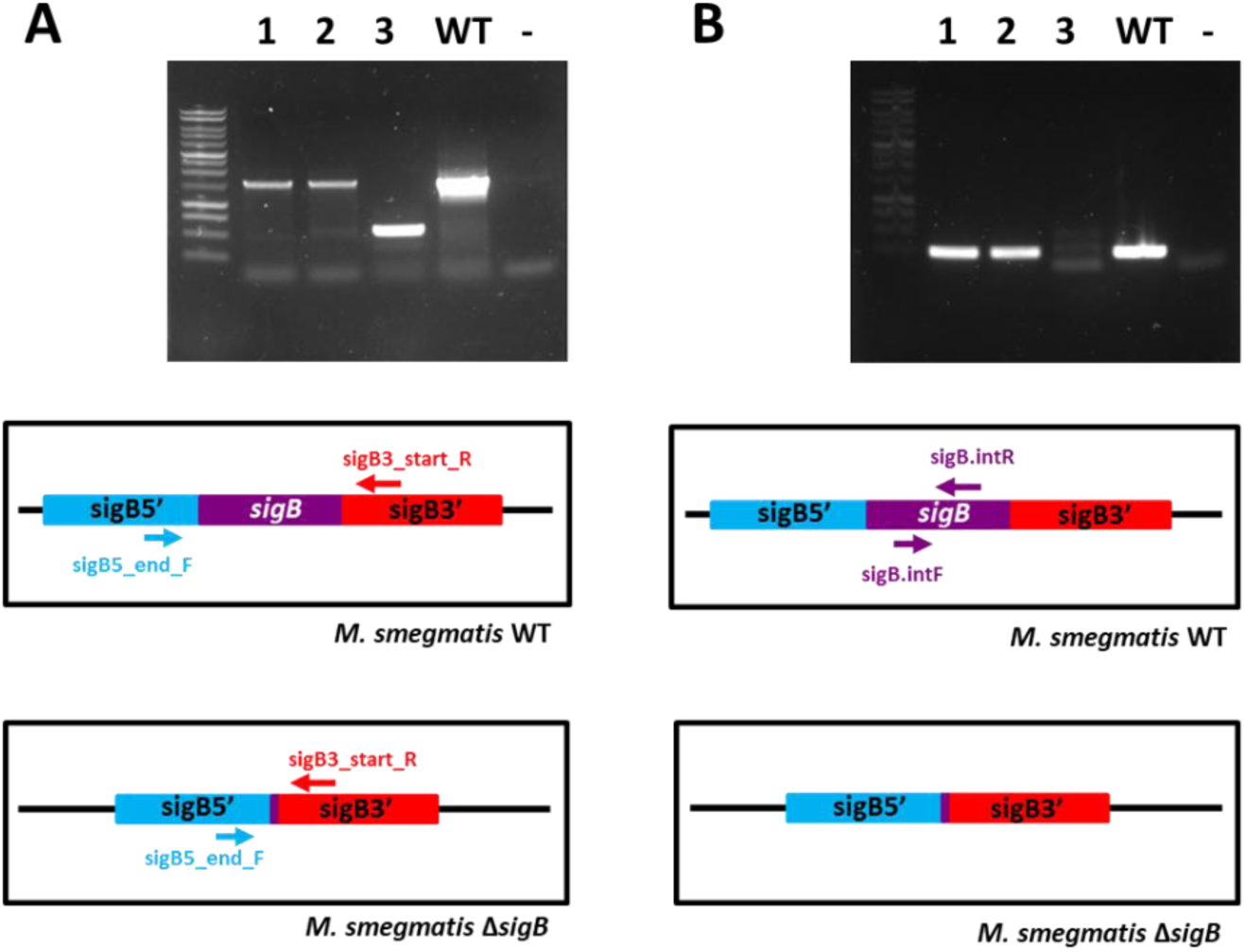
PCR verification results of the mutant. A. PCR performed with primers that hybridize to the sequences flanking the *sigB* gene. Expected band size for WT: 1431 bp. Expected band size for Δ*sigB*: 520 bp. B. PCR performed with primers that hybridize to the internal sequence of *sigB*. Expected band size for WT: 263 bp. Expected band size for Δ*sigB*: absence of a band or nonspecific amplification (as seen in lane 3). The sequence of the oligonucleotides is provided in **Table S3**. As a positive control, a PCR was performed using genomic DNA from *M. smegmatis* (WT) as the template. As a negative control (−), a PCR was performed without template DNA. The gel shows the 1 kb DNA marker (Promega). Below each gel, the location where the primers hybridize in the genome of a WT strain and a Δ*sigB* strain is depicted. Based on the gel observations, it was concluded that colonies 1 and 2 have the genotype of a WT strain, while colony 3 has the genotype of a Δ*sigB* mutant strain.

